# Trends in stomatal density and size in maize hybrids representing 100 years of long-term breeding for yield

**DOI:** 10.1101/2025.10.21.683719

**Authors:** Memiş Bilgici, Elnaz Ebrahimi, Leticia Prada de Miranda, Sara Lira, Lucas Borras, Thomas Young, Recep Yavuz, Kenneth J Moore, Philip Dixon, Thomas Lübberstedt

## Abstract

Maize hybrid breeding started over 100 years ago, has increased yield and vigor through improved genetics in conjunction with increased fertilizer and pesticide use, planting density, and agricultural mechanization. Stomata are expected to change in response to rising atmospheric CO_2_ concentration and average temperature anomalies (°C). Yet, the impact of long-term maize breeding over the past century on stomatal traits and their responses to climate factors remains poorly understood. We evaluated stomatal traits at the seedling stage in 27 maize hybrids released from 1920 to 2022, grown under controlled conditions. Modern hybrids (2013 ‒ 2022) had a smaller total stomatal pore area (9.17 x 10^8^ μm^2^) than (1920 ‒ 2012) historical ERA hybrids (9.94 x 10^8^ μm^2^; *p*≤ 0.001), a higher stomatal density (47.2 per mm^-2^) vs. historical ERA hybrids (44.5), and a smaller leaf area (17.9 cm^2^ vs. 20.5 cm^2^). No significant differences were found in the size (μm^2^), length (μm), or width (μm) of stomata between the two groups. Stomatal density increased, while all other traits decreased in modern hybrids. Stomatal density was negatively correlated with stomatal size (r = -0.62), length (r = -0.57), width (r = - 0.54), and leaf area (r = -0.54). Stomatal size had a negative correlation with atmospheric CO_2_ concentration (r = -0.22) and average temperature anomalies (°C) (r = -0.35) in the hybrid’s year of release and climate proxies. In contrast, stomatal density had a positive correlation with both atmospheric CO_2_ concentration and average temperature anomalies (°C) (r = 0.44) in year of release. Total stomatal pore area negatively correlated with atmospheric CO_2_ concentration (r = -0.45) and average temperature anomalies (°C) (r = -0.36). Our study indicates that maize stomatal traits suggest inadvertent selection for key stomatal traits (density and size), total stomatal pore area (per year decline of about 0.02%) associated with yield stability and environmental adaptation.

**Highlights:**

Maize stomatal traits changed through environmental (CO_2_ and °C) adaptation but total stomatal pore area, effects indirectly by decreased leaf area in maize hybrids representing 100 years of long-term breeding for yield.
A negative correlation was observed across 27 ERA hybrids between stomatal density and stomatal size, length, width and leaf area. Stomatal density increases while stomata size, length width and leaf area decrease per se.
Over the past 100 years, the total stomatal pore area on leaves decreased, while stomatal density increased as leaf area declined, revealing a connection between these two patterns.
A negative correlation was found between total stomatal pore area and atmospheric CO_2_ concentrations, and temperature over the past century.

## Introduction

Maize hybrid breeding started over 100 years ago (Shull, 1908; East, 1908) and resulted in significant yield increases. Double-cross hybrids (crosses between two single-cross hybrids) made maize hybrids economically viable (Jones, 1922). In 1926, Henry A. Wallace founded Hi-Bred Corn (later known as Pioneer Hi-bred), the first company to commercially produce and sell hybrid maize seeds. It is still one of the leading companies today (as part of Corteva Agriscience). Currently, the majority of maize varieties in the U.S. and worldwide are hybrids. In the U.S., average maize yield has increased 7-fold since the early 1900s, rising from about 1.4 Mg ha^-1^ to 11.3 Mg ha^-1^ in 2023 (Duvick, 2005; Nielsen, 2023; USDA 2025).

Changes in agriculture significantly shaped the development of modern hybrids over the past 100 years, such as the use of fertilizers and pesticides, and mechanization. Key drivers of yield gains in modern hybrids include tolerance to higher planting densities; (Assefa *et al.*, 2018) and resistance to drought and lodging (Campos *et al.*, 2006; Cooper *et al.*, 2014), which led to changes in plant stature (Hammer *et al.*, 2009; Borrás and Vitantonio-Mazzini, 2018; Wang *et al.*, 2020; Ordóñez *et al.*, 2024). Most recently, breeding programs have focused on increasing harvest index (Ruiz *et al.*, 2023; Shao *et al.*, 2024), leading to the development of short-stature maize varieties capable of tolerating even higher planting densities (Kosola *et al.*, 2023). In contrast, the potential yield per plant has not changed (Tokatlidis and Koutroubas, 2004; Duvick, 2005; Lee and Tollenaar, 2007). These advances have emerged during the period of rapidly changing climate conditions.

During the last century, atmospheric CO_2_ levels significantly increased from around 280 to 420 parts per million (ppm) (Tans and Keeling, 2016; Lan *et al.*, 2024). The Mauna Loa Observatory in Hawaii recorded CO_2_ levels of 428 ppm in March 2025 (https://gml.noaa.gov). If current emission trends continue, CO_2_ concentrations are projected to exceed 540 ppm by the year 2100 (Meinshausen *et al.*, 2011; Cui *et al.*, 2024). Global surface temperatures have increased by over 1 degree Celsius compared to the late 19^th^ century pre-industrial average which is also accompanied by frequent extreme weather events (Coumou and Rahmstorf, 2012; Seneviratne *et al.*, 2021; IPCC, 2022 and 2023). These environmental changes pose a challenge to maize productivity (Gong *et al.*, 2015; Ainsworth and Long, 2021; Rezaei *et al.*, 2023). Maize yield forecasts have become considerably more pessimistic (Li *et al.*, 2022). The increased frequency of droughts, floods, and storms can severely damage crops (Webber *et al.*, 2018). Changes in rainfall distribution can result in water stress (Kim and Lee, 2023), while higher temperatures (Lesk *et al.*, 2022) can accelerate plant development, shorten the growing season, leading to lower yields (Jägermeyr *et al.*, 2021).

In maize, a C₄ plant, mesophyll cells capture CO_2_ and incorporate it into four-carbon compounds, which are transported to bundle-sheath cells. There, CO_2_ is released for assimilation by RuBisCO in the Calvin cycle, enhancing photosynthesis, especially under high temperatures and light conditions (Zafar and Bailey-Serres, 2025). Stomata, microscopic pores on lower leaf surfaces, are important for gas exchange and thus directly connected with multifactorial climate change (Yan *et al.*, 2017; Hepworth *et al.*, 2018; Haworth *et al.*, 2023; Gray and Dunn, 2024). By opening, they enable CO_2_ uptake for photosynthesis and drive transpiration, which maintains water and nutrient flow from roots to shoots. Under limited soil moisture, evaporative demand can exceed the plant’s capacity to supply water. Leaf water potential declines, turgor is lost, and stomata begin to close, increasing stomatal resistance (rₛ) to CO_2_ diffusion. This closure conserves water but restricts CO_2_ uptake, reducing photosynthesis, carbon gain, and growth (Sage, 2004; Lawson and Blatt, 2014; Buckley, 2019; Bouvier *et al.*, 2024). Both stomata pore area and duration of opening are key determinants of plant’s water-use efficiency (WUE) (Dittberner *et al.*, 2018; Henry *et al.*, 2019; Barl *et al.*, 2025; Singh *et al.*, 2025; Zhang *et al.*, 2025).

The unique Pioneer ERA hybrids span 100 years of maize hybrid breeding and offer an opportunity to study changes in stomatal traits over this period. ERA studies have focused on overall yield improvement, and correlated traits, such as genetic gain, high planting density (Duvick, 2005), nitrogen (Haegele *et al.*, 2013; York *et al.*, 2015; DeBruin *et al.*, 2017; Mueller *et al.*, 2019), water and radiation use efficiency (Messina *et al.*, 2022), and harvest index (Campos *et al.*, 2006; Cooper *et al.*, 2014; Kosola *et al.*, 2023). Duvick and Cassman (1999) linked historical yield gains to reduction in tassel size and increases in stay-green traits. Campos *et al*. (2006) and Reyes *et al*. (2015) demonstrated that modern hybrids have shorter anthesis-silking intervals and experience less barrenness under drought stress (Smith *et al.*, 2014). Over the past eight decades of breeding, Pioneer maize hybrids developed smaller root systems, which are likely to facilitate higher plant population densities (Rinehart *et al.*, 2024), but a water capture capability similar to older varieties with larger root systems (Messina *et al.*, 2022). Perez *et al*. (2019) found that European hybrids exhibited reduced early leaf area but little change later in the growth period. Overall, gains in leaf area index were primarily driven by increased planting density, while hybrid improvements enhanced density tolerance (Kalogeropoulos *et al.*, 2024). The average leaf angle changed as the leaves became more erect (Elli *et al.*, 2023). Maize water-use efficiency and harvest index have increased over the past decades (Rotundo *et al.*, 2025). While ERA hybrids have been intensely studied for agronomic traits, little is known regarding changes at the cellular level.

This study quantifies century-long changes in maize stomatal traits. We investigated stomatal traits in maize seedlings of 27 hybrids grown under the controlled conditions. We evaluated stomata length, width, size, total stomatal pore-to-leaf area, and leaf area for 27 maize hybrids released between 1920 and 2022. Our objectives were to (a) quantify whether and how stomatal traits were changing over the past century, (b) assess the relationship between stomatal traits and changing environmental conditions in the past 100 years, and (c) discuss implications of stomatal trait changes for guiding future maize breeding strategies.

## 2. Materials and Methods

### 2.1. Plant Materials

We used a total of 27 ERA hybrids representing the period of 1920 to 2022 including 14 historic ERA hybrids (1920-2011) and 13 modern hybrids (2013-2022). Hybrid seeds (Table 1) were provided by Corteva Agriscience.

**Table 1.** Maize hybrids used in this study, their year of release, and corresponding ERA Historic and Modern.

| Hybrid identification code in the study | Hybrid Name | Year of commercial release |
| --- | --- | --- |
| **Historic ERA** |  |  |
| PHI01 | Reid | 1920 |
| PHI02 | 307 | 1936 |
| PHI03 | 352 | 1946 |
| PHI04 | 354 | 1953 |
| PHI05 | 354A | 1958 |
| PHI06 | 3376 | 1965 |
| PHI07 | 3366 | 1972 |
| PHI08 | 3382 | 1976 |
| PHI09 | 3378 | 1983 |
| PHI10 | 3379 | 1988 |
| PHI11 | 3394 | 1991 |
| PHI12 | 33P67 | 1999 |
| PHI13 | 33T59 | 2007 |
| PHI14 | P1151 | 2011 |
| **Modern (still in use or marketed)** |  |  |
| PHI15 | P1197 | 2013 |
| PHI16 | P1366 | 2017 |
| PHI17 | P0574 | 2014 |
| PHI18 | P1197 | 2014 |
| PHI19 | P1185 | 2015 |
| PHI20 | P0953 | 2021 |
| PHI21 | P0995 | 2021 |
| PHI22 | P1082 | 2021 |
| PHI23 | P1222 | 2021 |
| PHI24 | P1244 | 2018 |
| PHI25 | P1413 | 2022 |
| PHI26 | P1548 | 2020 |
| PHI27 | P1587 | 2020 |

### 2.2. Growth Chamber Experiments

A growth chamber (CONVIRON Flex; 3.6 m^2^ growth area, 635 mm growth height) is equipped with light-emitting diodes (LEDs) to deliver a light intensity of 1000 µmol m^-2^ s^-1^ at 152 mm from the light source. Light output was monitored with a Quantum Light Meter (Apogee Instruments, Logan, Utah, USA). A photosynthetic photon flux density (PPFD) of 742.8 µmol m^-2^ s^-1^ within the photosynthetically active radiation (PAR) range of 400-700 nm was applied to the 27 maize hybrids (1920-2022). Photon flux densities were measured in the same growth chamber for 3 runs as follows: 0.83 µmol m^-2^ s^-1^ in the ultraviolet (UV) range, 124.3 µmol m^-2^ s^-1^ (400-500 nm), 294.7 µmol m^-2^ s^-1^(500-600 nm), 323.8 µmol m^-2^ s^-1^(600-700 nm), and 159.1 µmol m^-2^ s^-1^ (700-780 nm).

The daytime and night-time temperatures were maintained at 25°C and 15°C, respectively, with a light cycle of 14 hours of light and 10 hours of darkness and a relative humidity of 65% ± 5%. The chamber was regularly monitored to maintain proper temperature, humidity, and light intensity, ensuring plants remained healthy and free from stress or disease. All plants received the same amount of tap water, applied as needed to keep the soil consistently moist without waterlogging.

For each of 27 hybrids, nine plants were evaluated per experiment (Averaged the three plants to obtain one block-tray experimental unit to avoid pseudo replication at the plant level). The study was repeated in three independent growth chamber experiments, each using a randomized complete block design with three replications (27 genotypes × 3 plants as smallest experimental unit (“plot”) in each block × 3 replicates). To minimize the effects of micro-environments, the positions of 3-plant entries in trays were randomized within each run. Planting trays (15-cell trays, 127 mm deep) were filled with moistened Berger BM7 bark mix soil containing a starter fertilizer, bark, coarse perlite, dolomitic and calcitic limestone, premium coarse peat moss, and a non-ionic wetting agent. Maize seeds were planted at a depth of approximately 20 mm.

### 2.3. Plant measurements and stomatal imaging

Plant measurements were taken 14 days after planting across all hybrids, when the seedlings reached the four to five leaf stages (V4-V5). Because emergence occurred at different times-unevenly across plants within genotypes and replications, the growth stage was used as a reference for measurements (14 days). The second leaf (continuing from the base) was selected for both leaf area (LA) and stomatal trait (size, density, length, and width) assessments. LA was calculated as *leaf length x leaf width x 0.75* (Elings, 2000), where width was recorded at the widest point of the blade, and length measured from the base of the leaf blade to the tip.

For stomatal measurements, a thin layer of clear nail polish was applied to the abaxial (lower) epidermis at the widest point of the blade and allowed to dry completely. A strip of Scotch® Transparent Tape (3M, MN, USA) was then pressed onto the dried area and peeled away, lifting a cast of the epidermis containing stomatal imprints (Wu & Zhao, 2017). The tape was mounted on a glass slide for microscopic observation.

Microscopy was performed using an Olympus BX40 microscope equipped with a high-definition camera, trinocular viewing head, C-mount adapter, and 40 x / 0.65 Ph2 ¥/0.17 objective lens and 10x ocular lens oil-immersion objective. The microscope allowed real-time observation of stomatal impressions and facilitated image and video capture onto a removable SD card. Stomata impressions were imaged at 40 x magnification with a 10 x ocular lens using a Moticam S6 camera and Motic Analysis 3.1 software. High-resolution images were analyzed to quantify average stomatal size, pore area (µm^2^), length (µm), width (µm), and density (number of stomata per mm^2^). Stomatal density per LA was estimated using the size of the imaged field at the specified magnification and the measured leaf area. All stomata images from the all-hybrids dataset 1920 to 2022 were manually reviewed to eliminate false positives and false negatives. Pixel-based measurements were converted to micrometers (µm) using a predetermined scale factor derived from a calibrated micrometer slide. Final stomatal dimensions and densities were used for statistical analyses.

### 2.4. Total stomatal pore area calculation

The total stomatal pore area per leaf was estimated for each genotype as: total stomatal pore area (𝜇𝑚^2^ leaf^-1^) = leaf area (𝑚𝑚^2^) × stomatal density (stomata no. 𝑚𝑚^-2^) × stomatal size (pore area) (𝜇𝑚^2^ stomata^-1^).

In this study, stomatal density and average size, pore area were obtained from image analysis, and LA was measured on the second leaf (cm^2^), converted to *mm^2^* prior to calculation as described above. Because size was measured in *µm^2^*, the calculated total pore area is expressed in *µm^2^*. This calculation was used to evaluate changes in total pore area across hybrids released over the past century.

### 2.5. Data Annotation and Model Training

A total of 192 maize isogenic line pairs (Yavuz et al. 2025) were used to train a model designed to predict stomatal position, orientation, and size on unrelated datasets. Images were derived from tape-mounted stomatal impressions captured by microscope, as described above.

Stomata features in each image were manually annotated using rotated-rectangle bounding boxes created with custom Python software to identify each stomate. The YOLOv8 Object Bounding Box (OBB) model (Jocher *et al.*, 2023) was trained via transfer learning on annotated stomata dataset, employing data augmentation techniques such as rotations, translations, scaling, and mix up to enhance robustness. Its performance was evaluated using metrics such as precision, recall, and mean average precision (mAP). Training was conducted over 100 epoches. The learning rate was dynamically adjusted during training to optimize model performance. The model’s performance on validation set achieved a precision of 0.83, recall of 0.15, and mAP@50 of 0.31 (Figure 1) (Jocher *et al.*, 2023). Final box loss and classification loss were 0.78 and 0.45, respectively. Average Intersection over Union (IoU) for the validation set was 0.78. The average centroid distance was 14.1 pixels, and the average angle difference was 6.78 degrees.

The data were separated using the ultralytics.data.split_dota module (Jocher *et al.*, 2023) into 70% training, 15% validation, and 15% for testing. Climate data

We analyzed year of release climate proxies (annual-mean atmospheric CO_2_ and annual temperature anomaly) from 1920 to 2022 (Fig 9. S1). (1958-present from Mauna Loa record, for earlier years from Law Dome ice-core CO_2_, harmonized). The atmospheric CO_2_ data were obtained from the Mauna Loa Observatory recordings accessible from the Earth System Research Laboratory (Climate Change: Atmospheric Carbon Dioxide at www.esrl.noaa.gov and accessed on April 15, 2025). Temperature anomalies, were sources from NOAA’s Merged Land-Ocean Surface Temperature dataset, using annually mean values for the same period (Fig 10. S1; https://www.ncei.noaa.gov, accessed on April 15, 2025).

### 2.6. Statistical Data Analyses

All statistical analyses were conducted using R software 4.4.3 (R Core Team, 2024) with RStudio (RStudio Team, 2024) (http://www.r-project.org). Traits were regressed using linear mixed effect models, with year of release as a fixed effect and hybrids, rep (block) as a random effect to evaluate changes over the period of the released-breeding period. Using a mixed model approach, the stomatal traits of each hybrid were expressed as: model and error structure; 𝑖 =1,…27, 𝐻 index hybrids, 𝑟 = 1,2,3 index growth-chamber runs (experiments), and 𝑏 =1,2,3 index blocks within runs. We used the following model: 𝑌𝑖𝑟𝑏 = 𝑢 + 𝐻𝑖 + 𝐶𝑟 + (𝐻𝑥𝐶)𝑖𝑟 + 𝑅𝑏(𝑟) + 𝑒𝑖𝑟𝑏, where 𝑌𝑖𝑟𝑏 is the response for hybrid 𝑖 in block 𝑏 of run 𝑟 ; 𝜇 is the intercept; 𝐻𝑖 is the fixed hybrid effect; 𝐶𝑟 is the random effect of run; (𝐻𝑥𝐶)𝑖𝑟 is the random hybrid-by-run interaction; 𝑅𝑏(𝑟) is the random (rep) block effect nested within run; and 𝑒𝑖𝑟𝑏 is the residual. For tests of hybrid main effect, we used the 𝐻𝑥𝐶 interaction as the error term (denominator), which evaluates whether hybrid differences are consistent across replicated runs.

We analyzed the differences between historic hybrids (released from 1920 to 2012) and modern hybrids (released from 2013 to 2022). The significance of differences between means was assessed using two-sided Student’s t-tests at a 95% confidence level (α = 0.05). Pearson correlation coefficients were calculated to analyze relationships between the evaluated traits. Relationships between the year of release and traits were analyzed using linear least-squares regression. R^2^ calculations of marginal and conditional effects for the models were performed using the “performance” package (Lüdecke *et al.*, 2021).

Using the ‘lmer’ function from the ‘lme4’ linear random-effects models were employed to determine Best Linear Unbiased Predictors (BLUPs) for each hybrid’s stomatal traits. Broad-sense heritability (H^2^) was calculated using the formula 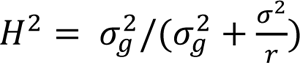, where; 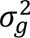 represents the genotypic variance, 𝜎^2^ is the error variance, and 𝑟 is the number of replications in the growth chambers per experiment. The calculation was performed using the lmer package in the *lme*4 R package (Bates *et al.*, 2015) and *nlme* (Pinheiro *et al.*, 2025). Figures were created for data visualization using the packages “ggplot2” (Wickham, 2016). Path analysis was performed to assess the total stomatal pore area using the ‘lavaan’ package (Rosseel, 2012). The proposed model incorporated both direct and indirect relationships among the variables.

Linear mixed-effects models were fitted for each stomatal trait with z-scored covariates. Model M1 (environment only-CO_2_ and temperature anomalies):

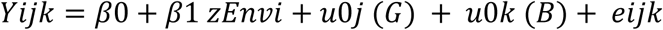

Model M2 (environment + leaf area):

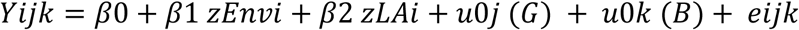

where u0j (G) ∼ N (0, 𝞼^2^G) is the random intercept for Genotype, u0k (B)∼ N (0, σ2B) is the random intercept for Block, and eijk ∼ N (0, σ2).

The environmental axis (zEnv) was the first principal component of standardized annual CO_2_ and temperature anomalies (both loadings = 0.707). To assess whether leaf area (explains away) the environment effect, we computed the attenuation metric of the zEnv slope from M1 to M2:

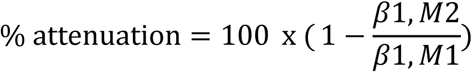

It quantified the relative contribution of zEnv vs xLA to fixed-effects R^2^ using Lindeman-Meranda-Gold (LMG) decomposition from an ordinary least squares model

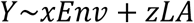

To test whether the effect of environment on stomatal traits is mediated by leaf area, fitted a simple path model observed zEnv as the exposure and leaf area as the mediator:

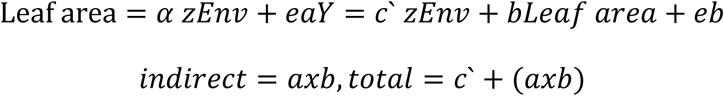

## 3. Results

### 3.1. Validation of measurement accuracy for stomatal traits

The stomata detection model, YOLOv8 OBB, demonstrated strong performance. The recall, a metric that measures the model’s ability to identify all positive instances in the dataset, was 97.9% (p ≤ 0.01). This indicates that the model correctly detected nearly all stomata present in the images, with very few cases missed. The precision was 86%, indicating that most of stomata identified by our model were correct although a small proportion were false positives. The F1 score was 91.6%. This indicates high reliability of our model for detecting stomata and avoiding misclassification. Mean Average Precision (mAP) is a reliable method for evaluating object detection models. It addresses challenges related to localization precision (IoU), confidence thresholds, and predictions across multiple classes. The mAP, calculated across IoU thresholds ranged from 0.50 to 0.95, is 53.5%. This indicates an expected decline in performance at stricter IoU thresholds, which is typical, as tighter matching presents greater challenges.

**Figure 1.**
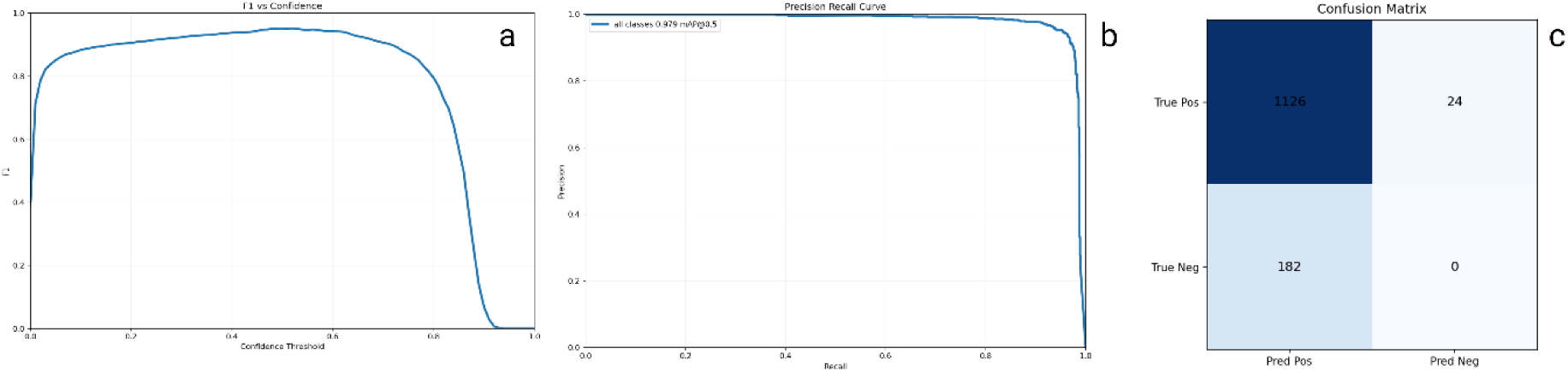
show a. F1 Score at Confidence Threshold Recall at Confidence Threshold, b. Precision at Confidence Threshold, and c. Recall at Confidence Threshold indicates the proportion of actual stomata correctly detected. A high recall rate indicates that there are few false negatives. F1 score is a harmonic mean of both precision and recall. It remains high only when both precision and recall are high. Average Precision at IoU Threshold of 0.50 metric. Average Precision at IoU Threshold of 0.50 metric shows the average precision when the Intersection over Union (IoU) threshold is set at 0.50. It implies that the model is considered correct if the predicted bounding box overlaps the true bounding box by at least 50%. mean average precision (mAP) evaluates model performance over a range of overlap thresholds, from loose (0.50) to strict (0.95) box matches in increments of 0.05. The moderate value reflects the expected decline in precision at stricter thresholds, where more exact bounding box alignment is required.

### 3.2. Stomatal trait distributions

Most stomatal traits followed near-normal distributions with minor skewness (Figure 2 in supplementary). Stomatal size and density exhibited slight deviations, while total stomatal pore area showed a strong right-skewed distribution. Leaf area and stomatal length and width were normally distributed, displaying tight, symmetric variation. These descriptive patterns provide context for comparing historic ERA and modern hybrids. Composite traits displayed different characteristics. Stomatal size trait showed moderate skewness of about 0.8, with a kurtosis value a slightly peaked distribution. Stomatal density spanned from 35 to 54 mm^2^, clustered around 42 to 50 mm^2^, exhibited a skewness of -0.5. The kurtosis was 3.0. Leaf area ranged from 14 to 19 cm^2^ and peaked around 18 to 21 cm^2^. It exhibited a right skewness of 0.6 with kurtosis indicating a slightly platykurtic distribution. Total stomatal pore area ranged from 7.4 x 10^8^ to 1.26 x 10^9^ *µm^2^*, showing a pronounced right skewness of 2.0 and high kurtosis, which points to a heavy right tail. Stomatal length ranged around 41 mm (from 38.9 to 45.5 mm) while the width was approximately 26.5 mm (from 25.3 to 28.3 mm). These traits displayed tight, symmetric distributions, with skewness values around 0 and kurtosis close to 3, indicating a normal distribution.

### 3.3. Comparison of historic and modern ERA hybrids for stomatal traits

Highly significant genetic variation was found for all stomatal traits across the 27 hybrids (Df = 26, *p* ≤ 0.001). Across all hybrids, stomatal size ranged from 1008.2 to 1269.7 μm^2^, with a mean of 1098.4 μm^2^. Stomatal density varied from 34.9 to 54.4 stomata/mm^2^, averaging 45.8 stomata/mm^2^. Stomatal length ranged from 38.9 to 45.5 μm, with a mean of 41.2 μm, while stomatal width was between 25.3 and 28.3 μm, averaging 26.5 μm. Total stomatal pore area ranged from 7.39 × 10^8^ to 12.60 × 10^8^ μm^2^, with a mean of 9.57 × 10^8^ μm^2^. Leaf area ranged from 14.0 to 29.0 cm^2^, with a mean of 19.2 cm^2^.

The modern hybrids (2013 to 2022) had significantly (*p* ≤ 0.001) increased stomatal densities (number per mm^2^), and decreased leaf areas (cm^2^), and total stomatal pore areas (µm^2^), compared to historic ERA hybrids (1920 to 2011) (Table 2). Modern hybrids exhibited higher stomatal densities (mean: 47.2 stomata per mm^-2^; range: 39.7–54.4) than historic ERA hybrids (mean: 44.5 stomata per mm^-2^; range: 34.9–51.8), but smaller leaf areas (mean: 17.9 cm^2^; range: 14.0–22.9) compared to historic ERA hybrids (mean: 20.5 cm^2^; range: 14.9–29.0). In addition, the total stomatal pore area was smaller in modern hybrids (mean: 9.17 × 10^8^ μm^2^; range: 7.45 × 10^8^–11.84 × 10^8^ μm^2^) than in historic ERA hybrids (mean: 9.94 × 10^8^ μm^2^; range: 7.39 × 10^8^–12.60 × 10^8^ μm^2^). No significant differences were found in the size (μm^2^), length (μm), or width (μm) of individual stomatal between the two groups.

**Table 2.**
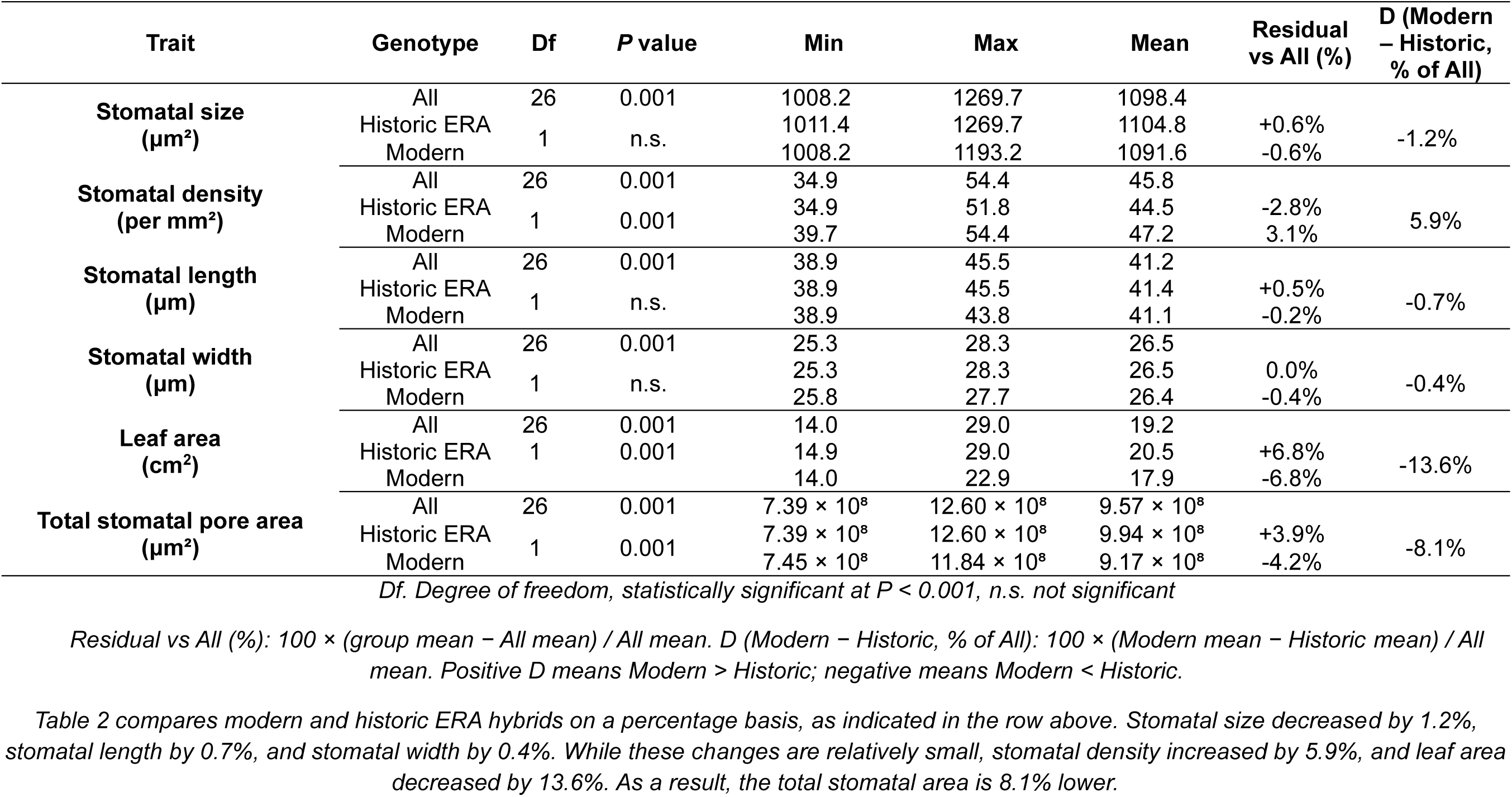
Comparative statistics for stomatal traits in modern (≥ 2013) and historic ERA hybrids.

**Figure 2.**
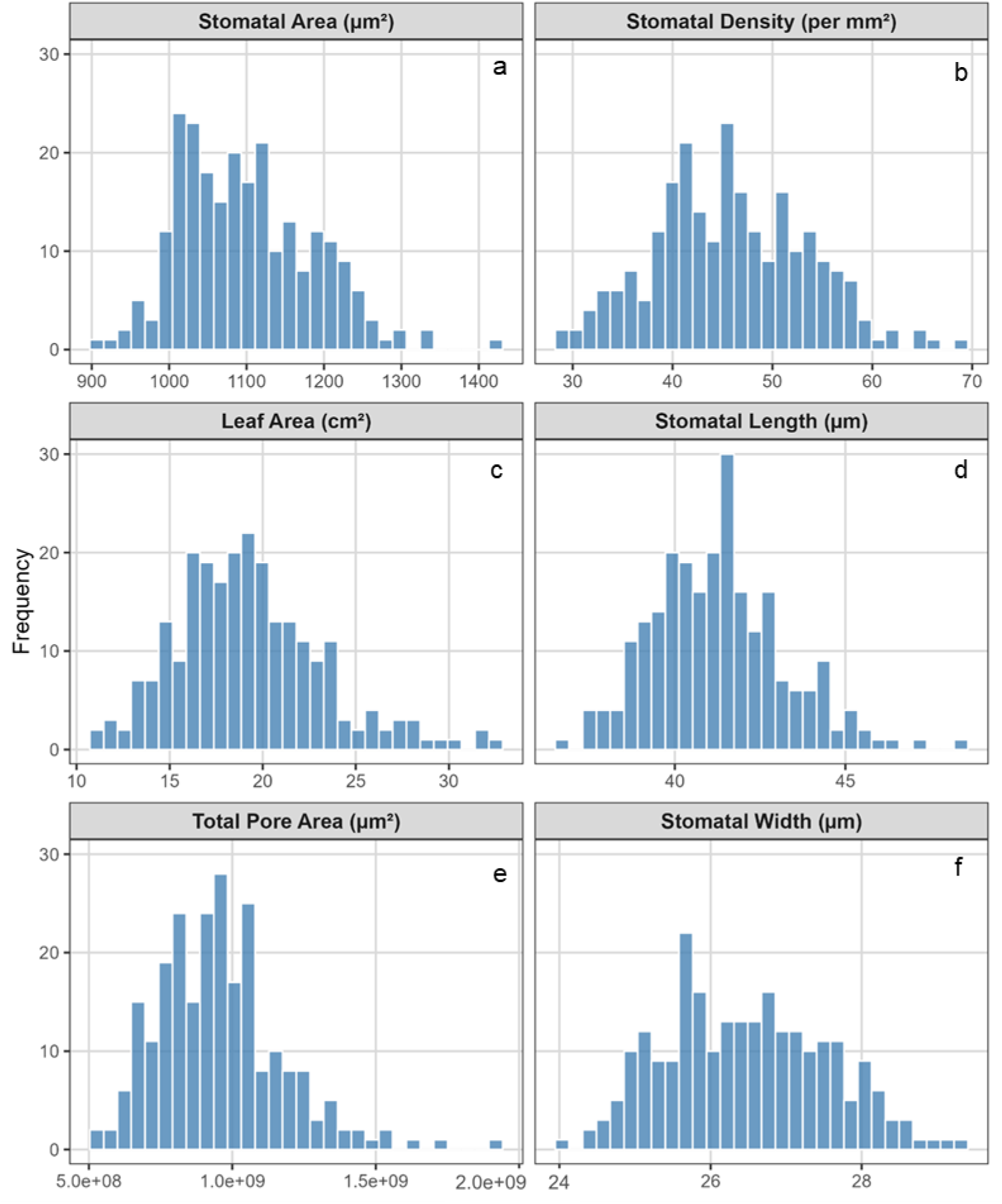
trait distributions across 27 hybrids: a. stomatal size (area); b. stomatal density; c. leaf area; d. stomatal length; e. total stomatal pore area; and f. stomatal width.

**Figure 3.**
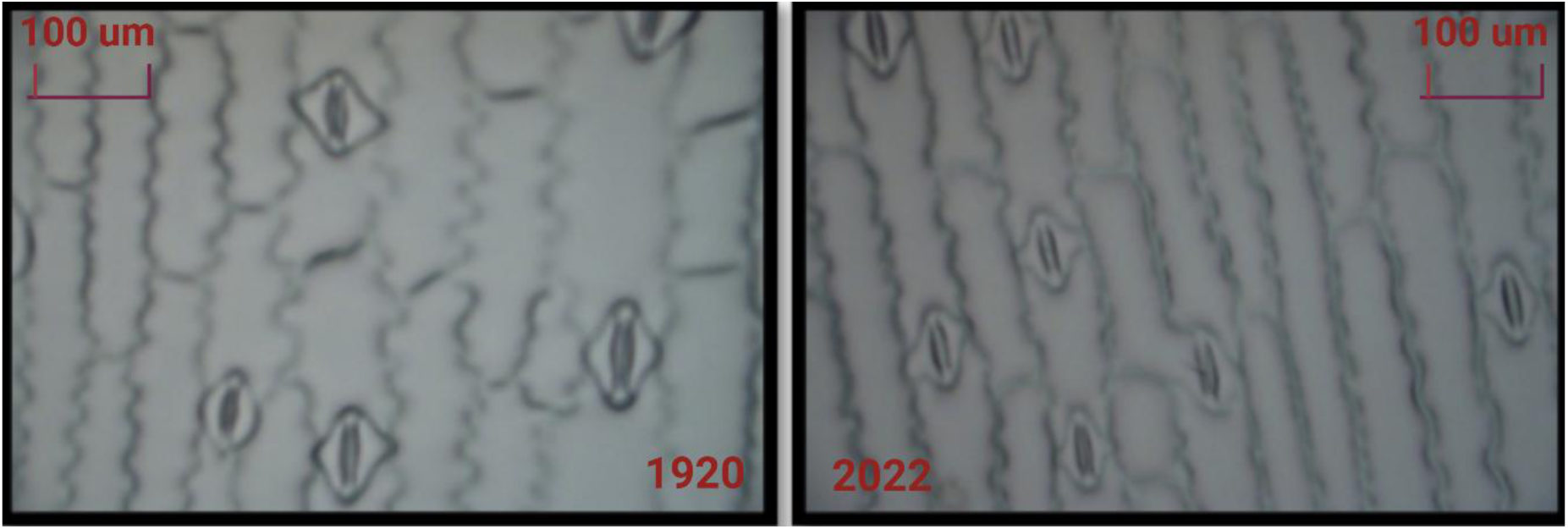
Microscopic images of stomata at 40X magnification from 1920 and 2022, indicating that the same magnification was used. Microns (µm) are employed to measure size standards

### 3.4. Heritabilities of stomatal traits

The broad sense heritabilities (H^2^) of all stomata traits ranged from 0.36 to 0.81 (Table 3). The highest heritability (0.81) with narrow confidence interval (CI 0.53–0.85) was found for total stomatal pore area, while stomatal density showed the lowest heritability of 0.36 (CI 0.12-0.61).

**Table 3.** broad-sense heritabilities (H^2^) for stomata traits.

| Trait | Broad Sense Heritability ($H^2$) | 95% Confidence Interval (CI)<br>Lower–Upper |
| --- | --- | --- |
| Leaf area ($\text{cm}^2$) | 0.74 | 0.60–0.88 |
| Stomatal size ($\mu\text{m}^2$) | 0.64 | 0.46–0.82 |
| Stomatal density (per $\text{mm}^2$) | 0.36 | 0.12–0.61 |
| Stomatal length ($\mu\text{m}$) | 0.68 | 0.51–0.85 |
| Stomatal width ($\mu\text{m}$) | 0.57 | 0.36–0.77 |
| Total stomatal pore area ($\mu\text{m}^2$) | 0.81 | 0.53–0.85 |

### 3.5. Trait correlations

**Figure 4.**
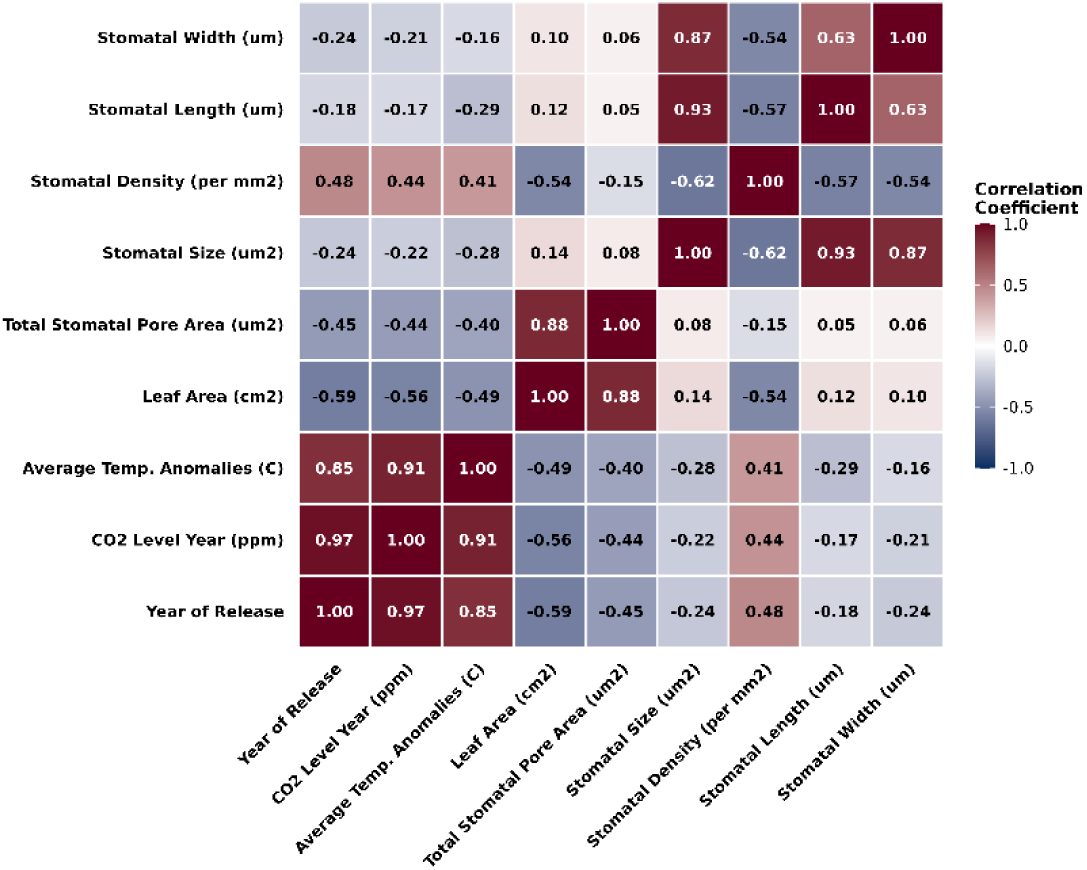
Pearson correlations were determined among all stomata and climate traits

Stomatal density was negatively correlated with stomatal traits such as stomatal size (r = -0.62), length (r = -0.57), width (r = -0.54) and leaf area (r = -0.54).

Stomatal size and length exhibited a strong positive correlation (r = 0.93), as stomata length increased, size also increased. Similarly, stomatal size and width showed a strong positive correlation (r = 0.87), which was expected since size was dependent on both dimensions. Stomatal length and width had a positive correlation (r = 0.63) stomata tended to grow proportionally in both dimensions.

Stomatal size showed a negative correlation of r= -0.22 with atmospheric CO_2_ concentration in parts per million (ppm) at the time the hybrid was released, whereas stomatal density exhibited a positive correlation of r= 0.44. Additionally, stomatal size had a negative correlation of r= -0.28 with average temperature, while stomatal density had a positive correlation of r= 0.41. Furthermore, the total stomatal pore area negatively correlated with both atmospheric CO_2_ concentration (r= -0.44) and average temperature anomalies (°C) (r= -0.40).

Rising CO_2_ levels and increasing temperatures effected stomatal traits. These findings demonstrated a clear response of stomatal traits to CO_2_ and temperature changed.

### 3.6. Path analysis

**Figure 5.**
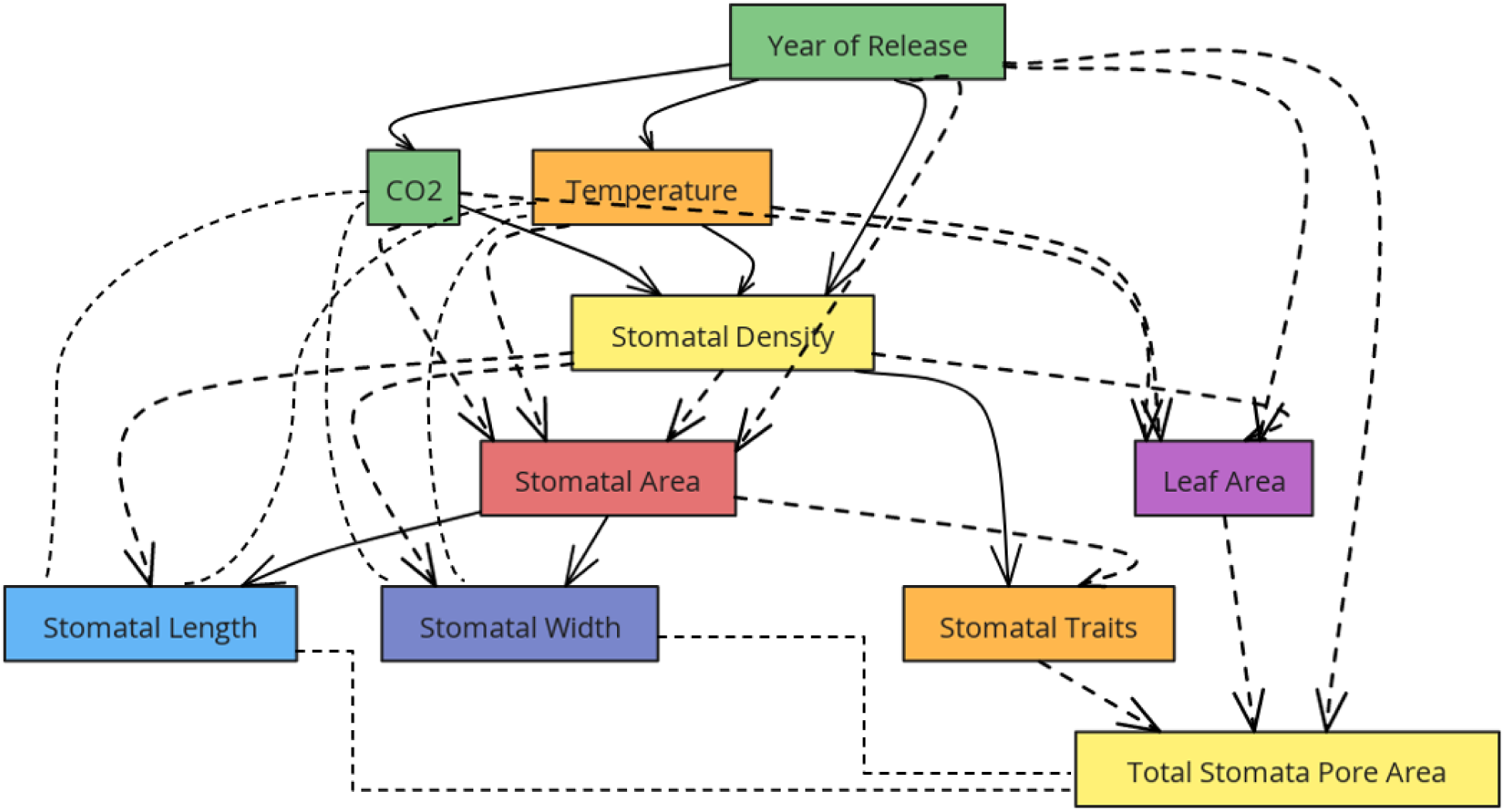
Path analysis for total stomatal pore area. Positive correlations are represented by solid arrows, while negative correlations are indicated by dotted-line arrows. Stomatal traits showed the effect of density and area in the legend.

Hypothesis 1: Stomatal traits directly and positively affect total stomatal pore area. Hypothesis 2: Leaf area indirectly affects total stomatal pore area through its influence on stomatal traits.

Hypothesis 3: Year of release, increased atmospheric CO_2_ concentration and temperature affect stomatal traits, leading to a reduction in total stomatal pore area.

Leaf area had the most significant impact on total stomatal pore area (d1=-0.93, *p*≤ 0.001). Stomatal density contributed significantly (d2=0.39, *p*≤ 0.001). Stomatal length had a slight negative effect (d3=-0.15, *p*≤ 0.05), while stomatal width had a modest positive effect (d4=0.17, *p*≤ 0.05). Increased CO_2_ resulted in widened and shortened pores while leaf size was reduced, so its net effect was close to zero. Stomatal length decreased with rising temperatures, while stomatal width remained unaffected.

Factors such as CO_2_ levels, temperature, and year primarily influenced total stomatal pore area through these mediators, as their direct effects were minimal. Increased CO_2_ and temperature exhibited a strong correlation (r=0.88, *p*≤ 0.001). CO_2_ was strongly negatively correlated with leaf area (r = −0.77), suggesting that higher CO_2_ levels may be associated with smaller leaves.

Temperature anomaly showed only a weak, nonsignificant positive link to leaf area (r=0.24). Larger leaves had lower stomatal densities but significantly greater total stomatal pore area (r=−0.57 and r=0.81, respectively). Temperature’s strongest indirect effect was via leaf area (+0.22), resulting in a total effect of (+0.13) on total stomatal pore area. The year of hybrid release had a net negative effect of (-0.18) because its pathway through leaf area (-0.73) outweighed the positive impact of stomatal density.

The overall effect of CO_2_ was negative (-0.39), mainly driven by the negative impact of stomatal density. Increased CO_2_ impacted leaf area, giving an indirect negative influence on total stomatal pore area.

H₁: Stomatal Traits Adaptation: Year → Stomatal traits. Predictions: The effects of year and environmental conditions persist even after controlling for leaf area. The partial correlations are approximately equal to the total correlations, and direct effects are detectable.

H₂: Leaf-Mediated By-Product via leaf area): Year → Leaf area → Stomatal traits (mediation). Predictions: The relationship from year to leaf area leads to stomatal traits through mediation. Adjusting for leaf area reduces the associations between year and stomatal traits, indicating that the path from year to leaf area is strong.

**Figure 6.**
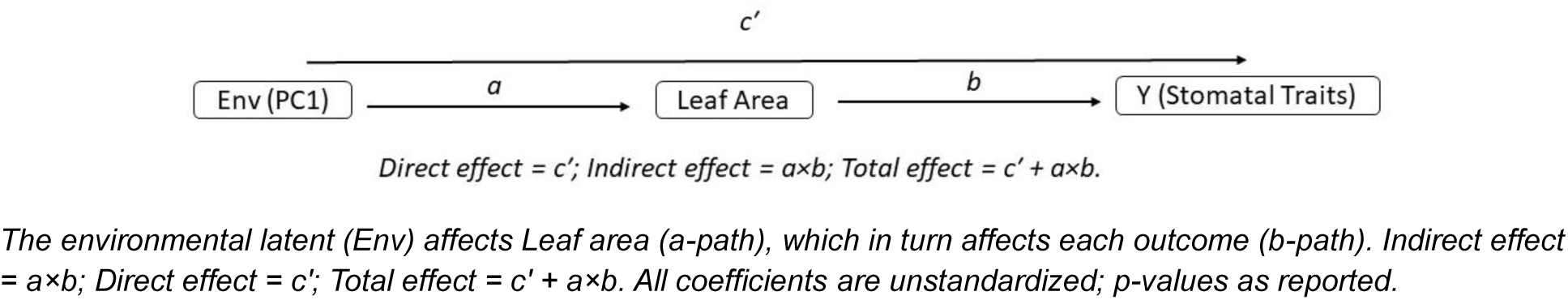
Structural equation model (SEM) depicting direct (c′) and indirect (a×b) pathways from the environmental axis (PC1) to stomatal traits via Leaf area.

**Table 4.** SEM summary including indirect and total effects.

| Trait | a (Env → LA) | b (LA → Trait) | Indirect | Direct (c') | Total Effect (c' + a×b) | Verdict |
| --- | --- | --- | --- | --- | --- | --- |
| Stomatal Density | -1.932<br>(p < .001) | -0.411<br>(p = .002) | 0.794 | 1.279<br>(p = .01) | 2.073 | Both direct and mediated |
| Stomatal Size | -1.932<br>(p < .001) | -0.187<br>(n.s.) | 0.362 | -17.23<br>(p = .006) | -16.87 | Direct only |
| Total Stomatal Pore Area | -1.932<br>(p < .001) | 39.8M<br>(p < .001) | -76.9M | 14.7M | -62.2M | Mediated |

Maize stomatal traits changed due to environmental (CO_2_ and °C) adaptation but total stomatal pore area, effects indirectly by decreased leaf area in maize hybrids representing 100 years of long-term breeding for yield. Leaf area was direct and indirect adaptation to environment changed.

### 3.7. Trends for Stomatal Traits Over Time

We observed that total stomatal pore area on the 2^nd^ leaf decreased with the year of release (Table 3). The trend from 1920 to 2022 (r= -0.45, r^2^= 0.20; p = 0.02), y = − 1992 x + 4.92 × 10^8^ also shows a significant decline. For each year, the total stomatal pore area decreased by approximately 1992 um^2^, which corresponds to an annual decline of about 0.02% relative to the initial mean value.

**Figure 7.**
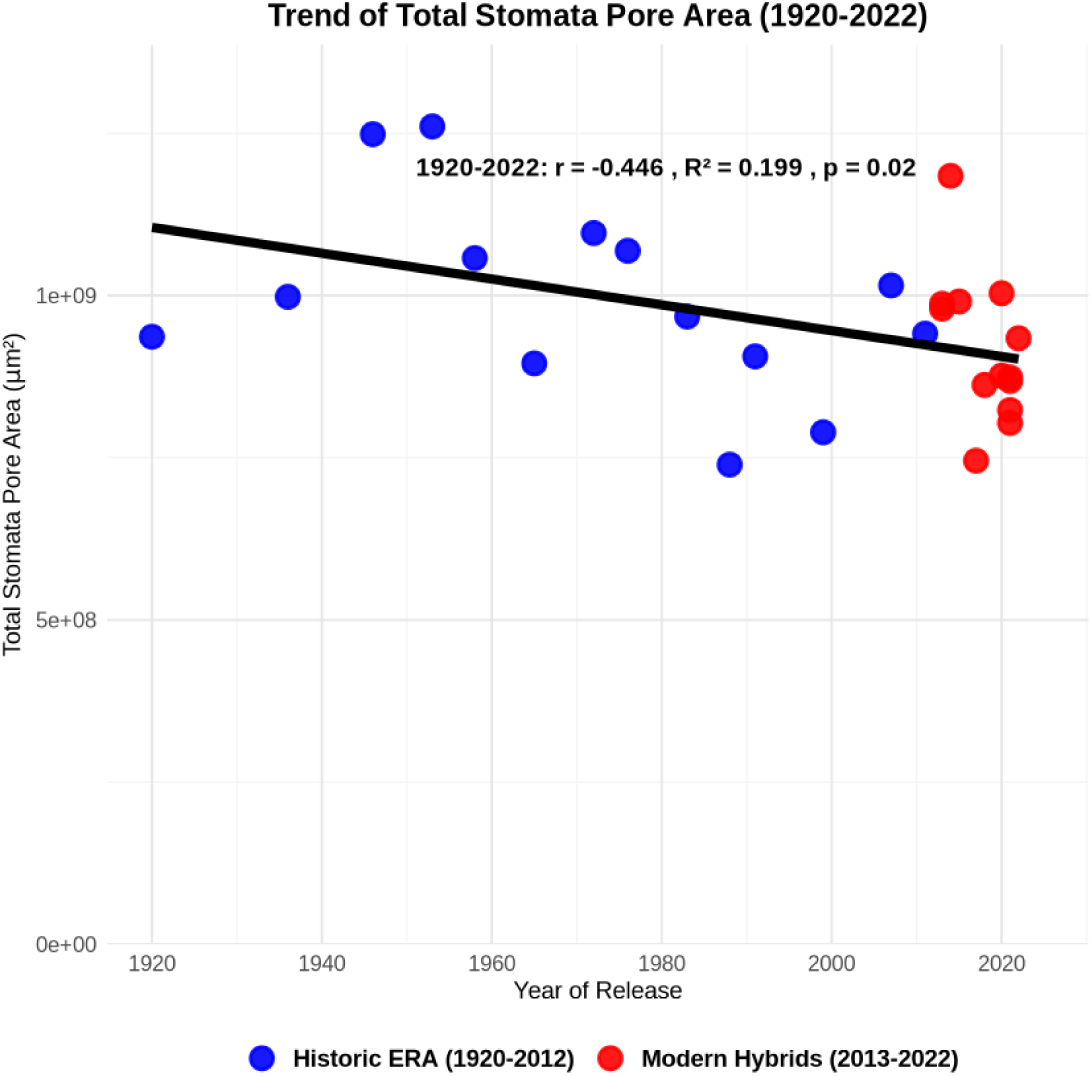
shows the trend of total stomatal pore area from 1920 to 2022.

From 1920 to 2022 distinct trends were observed for stomatal traits of 27 hybrids. Stomatal size shows a non-significant negative correlation (r = -0.25), while stomatal density increased significantly over this period (r = 0.46, *p*≤ 0.05), indicating an increase in the number of stomatal per unit area.

Conversely, the total stomatal pore area exhibited a significant negative correlation with the year of release (*p*≤ 0.05). This means that modern hybrids displayed a decrease in total stomatal pore area. Additionally, there is a strong negative correlation between stomatal size and density (r = -0.61, *p*≤ 0.05).

Modern hybrids have smaller stomatal size but a higher stomatal density compared to historic ERA hybrids. Thus, modern hybrids feature increased densities, but a smaller pore area than older hybrids. Moreover, plant leaf area decreased over the past 100 years.

We found a negative correlation between the total stomatal pore area of the entire leaf, CO_2_ levels (p= 0.017, r= -0.454, r^2^= 0.206) and temperature (r= -0.36). Additionally, stomatal density increased slightly with rising CO_2_ levels, with this relationship being statistically significant (*p*≤ 0.05). However, no significant relationship was observed between stomatal size and CO_2_ levels.

**Figure 8.**
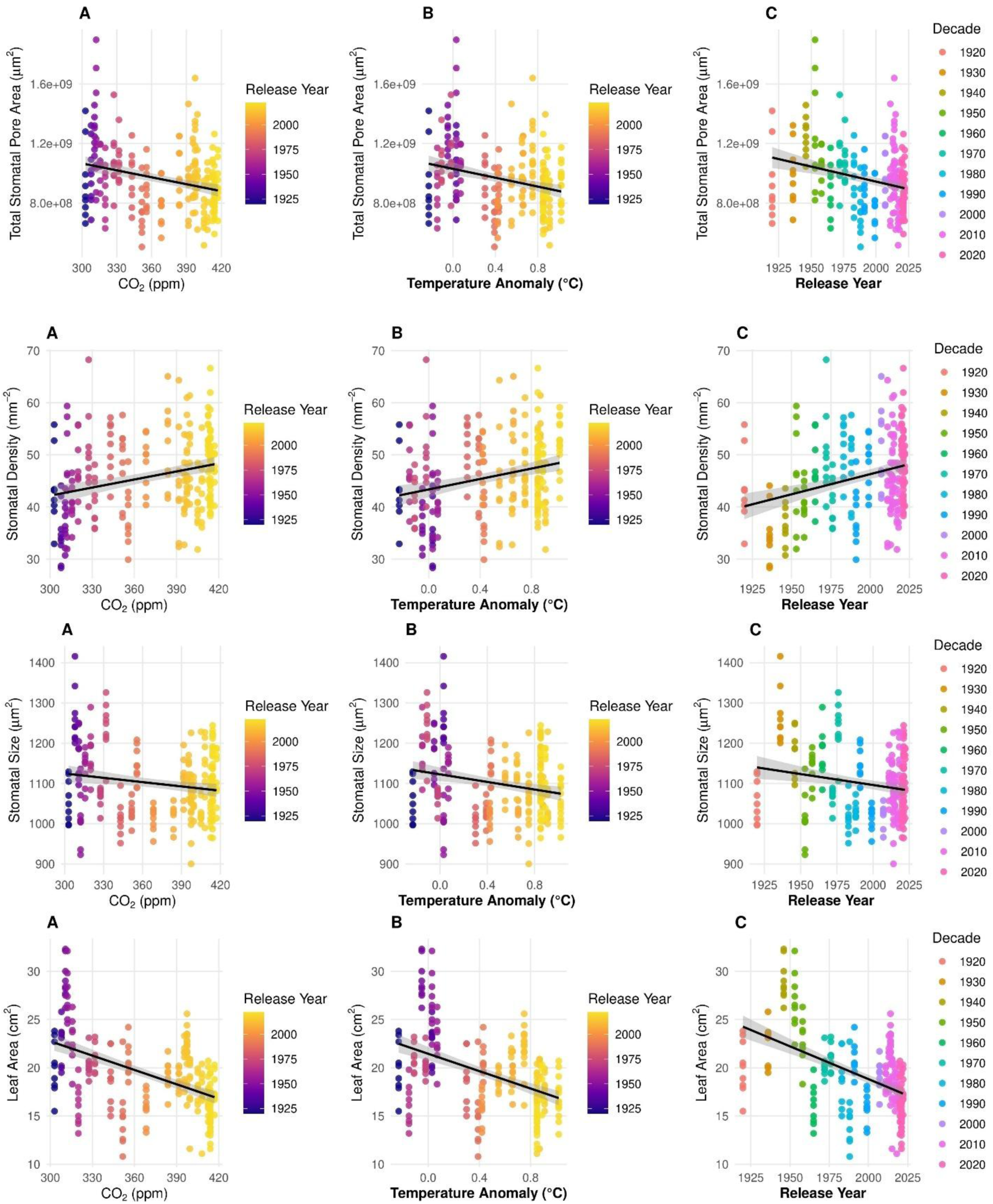
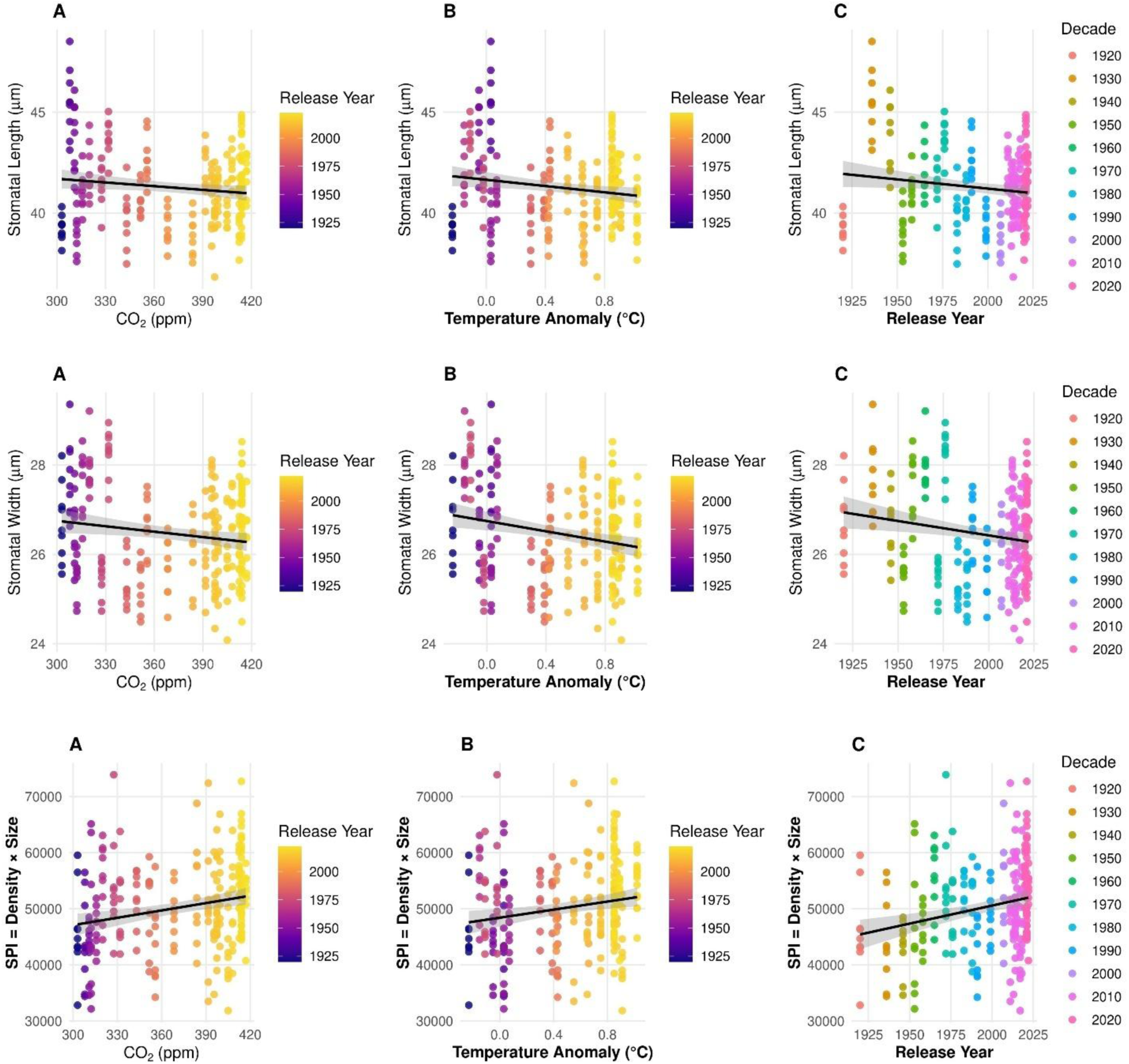
illustrates the total stomatal pore area, stomatal density, stomatal size, leaf area, stomatal length, stomatal width, and the interaction of stomatal density and size. This data is presented against three variables: (A) atmospheric CO_2_ levels (ppm), (B) mean temperature anomaly (°C), and (C) the year of release. The points represent replicate observations, which are colored according to the release year (in panels A and B) or by decade (in panel C). The lines depict linear regressions accompanied by 95% confidence intervals.

## 4. Discussion

The 27 hybrids used in our study are a small yet significant selection of maize germplasm, chosen for high agronomic importance in their respective eras between 1920 and 2022. In our study focusing on the 2^nd^ leaf, we assessed various stomatal traits. The broad sense heritability (H^2^) of these traits ranged from 0.36 to 0.81 (see Table 3). While this range is suitable for analysis under controlled conditions, it is not known whether stomatal traits of the 2^nd^ leaf are predictive for other leaves on the same plant. We conducted a field study where we collected stomata samples from the first leaf and the ear leaf of 10 different hybrids. While there was a significant difference between the first and ear leaf stomatal traits, they were closely correlated (r = 0.72). The measurements from the first leaf reliably predicted characteristics of the ear leaf (unpublished data). Long-term evolutionary changes in stomatal traits can be observed through field studies at varying CO_2_ concentrations. The study supports our findings, showing that stomatal density and number were low on new leaves but increased sharply during early development. Density stabilized once the leaves reached around 10 cm^2^, while the stomatal number continued to increase with growth and also depend on drought conditions (Zhao *et al.*, 2015). Importantly, stomatal traits at different developmental stages were not significant (Zhao *et al.*, 2015) and our finding closely correlated across genotypes (r=0.72) (unpublished data). This suggests that stomatal traits measured under controlled conditions for a defined leaf in a panel of genotypes can be predictive for stomatal traits of other leaves of these genotypes.

We observed changes in maize stomatal traits in leading 27 hybrids representing subsequent decades over the past 100 years. Modern hybrids had a lower total stomatal pore area than historic ERA hybrids. Modern hybrids tended to have denser but smaller stomata with a reduced total stomatal pore area. The trend towards hybrids with a reduced total pore area suggests stomata adaptation to ongoing changes in global climate, including rising atmospheric CO_2_ levels, temperatures, and intensified drought conditions.

A key question is whether increasing atmospheric CO_2_ concentration leads to a reduction in total stomatal pore area (increased stomatal density vs. decreased size) on maize leaves, as this would benefit the water balance of the plants. We observed changes in stomatal traits, including increased density (5.9%), decreased size (-1.2%) (not significant) and total stomatal pore area (-8.1%) per plant/leaf, compared to modern vs. historic ERA hybrids, resulting in an overall decline of about 0.02% per year over the past 100 years in the total stomatal pore area. Despite modern hybrids having a smaller total stomatal pore area per plant, they are grown at a higher planting density, which enables similar yields while absorbing more CO_2_ per area due to the increased total number of plant leaves (Kalogeropoulos **et al.*, 2024*). Over the past century, the CO_2_ level has increased by 1.5-fold (from 280 to 419 ppm). This implies that significantly more carbon was converted into grain yield, facilitated by the rise in CO_2_ concentration. Interestingly, Kalogeropoulos *et al*. (2024) found a decrease in the average leaf area of individual plants: a decline of 0.33% per year from 1983 to 2017 and by 6.7% from 1936 to 2014 (Rinehart *et al.*, 2024). Thus, although the average leaf area per plant has decreased (smaller leaves), the total leaf area per hectare has increased significantly. Today’s fields maintain or even exceed historic levels of total leaf area.

Our study demonstrated a close negative correlation between stomatal density and stomatal size (length and width), as well as leaf area in maize, aligning with short-term breeding studies (Zhao *et al.*, 2015; Serna, 2022; Mano *et al.*, 2023; Zhang *et al.*, 2025). Considering stomatal density, size, and leaf area together offers more insight than analyses of any single trait over the past 100 trends. Smaller stomata open and close more rapidly, improving control of gas exchange under water stress (Drake *et al.*, 2013; Chaves *et al.*, 2016; Liu *et al.*, 2019; Lawson and Vialet Chabrand, 2019; Clark *et al.*, 2022; Serna, 2022; Haworth *et al.*, 2023). Gas-diffusion theory indicates that small, widely spaced pores are disproportionately more efficient because diffusion scales better with pore circumference than area (Bidwell, 1974).

We hypothesized that increased atmospheric CO_2_ and temperature resulted in smaller, denser stomata. Indeed, total stomatal pore area per leaf was negatively correlated with CO_2_ and temperature in year of release, linking size, density, and leaf area to climate and grain yield (Woodward, 1987; Driscoll *et al.*, 2006; Ji *et al.*, 2008; Franks *et al.*, 2009; Yan *et al.*, 2017; Xu *et al.*, 2016; Liang *et al.*, 2023). Meta-analyses indicated that stomatal frequency generally declines as CO_2_ rises but can increase with higher temperatures and drought, depending on species and conditions. At higher CO_2_, stomata tend to be smaller and more densely packed (Yan *et al.*, 2017; Jordan *et al.*, 2020).

In maize, elevated CO_2_ (550 and 700 ppm) increased density but reduced size, conductance, and transpiration, improving WUE (Khan *et al.*, 2024; Sonmez *et al.*, 2023). These short-term studies support our long-term finding of increased density and decreased size, along with its adaptation to atmospheric CO_2_ concentration. However, these studies mentioned no yield gain benefit because C4 photosynthesis in maize is already near its CO_2_ efficiency maximum at 420 ppm, elevated CO_2_ does not directly stimulate photosynthesis or yield under non-stress conditions and projected benefits are mainly indirect (water-saving) under drought, with anticipated CO_2_-driven yield increases under dry conditions (Leakey *et al.*, 2006, 2009; van der Kooi *et al.*, 2016).

Some studies nonetheless report yield gains or heat-stress mitigation and higher WUE at elevated CO_2_, and increased temperature can reduce stomatal density (Abebe *et al.*, 2016; Lau *et al.*, 2018; Vajana *et al.*, 2024; Khan *et al.*, 2024; Bai *et al.*, 2025).

Our findings are consistent with other long-term adaptation studies in C3 plants. While only few long-term studies were conducted, investigation of an herbarium in Arabidopsis resulted in a century long-term study showing declining density with rising CO_2_ (Lang *et al.*, 2024). Similar long-term declines occurred in C3 angiosperm trees and ferns (e.g., *Pinus elliottii, Pinus taeda, Taxodium distichum*) over 150 years in Florida (Lammertsma *et al.*, 2011). A 40% density decrease across eight temperate tree species over 200 years accompanied a CO_2_ rise from 280 to 340 ppm and was supported by controlled experiments (Woodward, 1987). In contrast, our results in maize showed increasing stomatal density and supported adaptation to climate change such as increased atmospheric CO_2_ concentration and temperature, indicating that adaptive trends can diverge among species. Fossil and contemporary studies likewise associate density decline with rising CO_2_ (Hetherington and Woodward., 2003; Gienapp *et al.*, 2008; Doheny-Adams *et al.*, 2012; Vile *et al.*, 2012; Zheng *et al.*, 2013; Chen *et al.*, 2024; Rovira *et al.*, 2024; Klein *et al.*, 2025). C3–C4 physiological differences may explain those differences patterns. C4 maize generally uses water more efficiently and regulates stomata rapidly under hot, dry conditions, whereas C3 plants thrive in cool, moist environments but are more sensitive to water loss (Ozeki *et al.*, 2022; Way, 2012; Bertolino *et al.*, 2019; Song *et al.*, 2023). While long-term C3 crop studies are very limited, short-term studies in wheat and soybean showed that density can fluctuate with drought and heat (Chua and Lau *et al.*, 2024). A previous maize study involving 66 commercial hybrids released in Europe from 1950 to 2015 reported no changes in stomatal conductance or drought sensitivity, suggesting limited selective pressure on these traits over recent decades (Welcker *et al.*, 2022). However, stomatal traits were not measured.

This ongoing adaptation not only enhances maize grain yields but also plays a crucial role in addressing water conservation challenges in agriculture. Since 1950, rainfed maize yields in the USA corn belt have more than tripled without an increase in water inputs. The study revealed that the use of 61 ERA hybrids since 1934, has led to an increase in water use efficiency of 4.2% per year. This improvement is attributed to higher biomass productivity and better harvest index, with genetic advancements contributing 1.9% per year (Rotundo *et al.*, 2025). These findings are consistent with our observation that modern hybrids had a smaller average stomatal pore area, which could contribute to their improved water use efficiency.

Over the past 100 years, significant changes have occurred in global climate conditions, plant density, water use efficiency, and maize yield. However, the per-plant grain yield remained relatively unchanged (Tokatlidis and Koutroubas, 2004; Duvick, 2005). We observed changes in stomatal traits, including an increase in stomatal density, a decrease in stomatal size, and consequently, a reduction in the total stomatal pore area per plant or leaf. These changes in stomatal traits appear to be a consequence of indirect selection for yield and are correlated with increasing atmospheric CO_2_ concentration levels and temperature. We propose that selection for yield stability, drought resilience, and tolerance to higher planting densities have inadvertently shaped stomatal traits over generations under climate change environments. Over the past century, maize stomatal traits evolved through inadvertent selection associated with yield improvement and environmental adaptation.

Our findings showed that these changes generally favored increased stomatal density, reduced stomatal size, and smaller leaf area. These trends align with the biological expectation of a negative correlation between stomatal size and density, which likely results from geometric constraints and competition for epidermal space with other cell types, such as trichomes and glands (de Boer *et al.*, 2016; Nunes *et al.*, 2023; Chua and Lau, 2024). Based on our results, we propose that future maize breeding should prioritize smaller but more numerous stomata (density) per unit leaf area, which allows rapid adjustment of conductance and improved water-use efficiency. Efficient guard cell signaling, enabling quick responses to fluctuating conditions and light and vapor pressure deficit. These traits would not only improve resilience to climate change but also support higher yields. Intentional selection for stomatal traits offers a strategic advantage over inadvertent selection, particularly in drought-prone regions where water-use efficiency is critical.

## Acknowledgements

We thank Dr. D. Raj Raman for his comments regarding data interpretation. We also thank Diego Goncalves Caixeta, Kanogporn Khammona and Jianru Chen for their help collecting data. We would like to thank Corteva Agriscience for providing seeds.

This research was supported by Regenerating America’s Working Landscapes to Enhance Natural Resources and Public Goods through Perennial Groundcover (RegenPGC) is supported by Agriculture and Food Research Initiative Competitive Grant No. 2021-68012-35923 from the USDA National Institute of Food and Agriculture. Any opinions, findings, conclusions, or recommendations expressed in this publication are those of the author(s) and do not necessarily reflect the view of the U.S. Department of Agriculture.

## Data Availability Statement

The data supporting the conclusions of this article are included in the article and its supplementary data published online. The authors will make the raw data available upon request.

## Author contributions

MB: conceived the idea and research, methodology, performed experiments and analyzed data, literature search, original manuscript preparation; LP: performed experiments measurements, edited and reviewed the manuscript; EE, SL, LB, TY, RY, KJM, PD: wrote, edited and reviewed the manuscript; and TL: supervision, funding acquisition, original manuscript preparation, edited and reviewed. All authors wrote and reviewed the manuscript.

## Additional Supplementary

The Average of Each Hybrid on Measured Traits of the Second Leaf After 14 Days.

| Table 4. The Average of Each Hybrid on Measured Traits of the Second Leaf After 14 Days. |  |  |  |  |  |  |
| --- | --- | --- | --- | --- | --- | --- |
|  |  | Stomatal Density | Stomatal size-area | Stomatal length | Stomatal width | Total stomatal pore area |
| Release year | Genotype (hybrid) | Stomata mm<sup>-2</sup> | µm<sup>2</sup> | µm | µm | µm<sup>2</sup> |
| 1920 | Reid | 43.48 | 1056.85 | 39.26 | 26.70 | 9.36 × 10<sup>8</sup> |
| 1936 | 307 | 34.89 | 1269.65 | 45.45 | 27.71 | 9.97 × 10<sup>8</sup> |
| 1946 | 352 | 36.91 | 1171.23 | 43.68 | 26.63 | 12.48 × 10<sup>8</sup> |
| 1953 | 354 | 45.76 | 1011.38 | 39.17 | 25.54 | 12.60 × 10<sup>8</sup> |
| 1958 | 354A | 40.70 | 1139.09 | 41.20 | 27.52 | 10.57 × 10<sup>8</sup> |
| 1965 | 3376 | 47.53 | 1177.94 | 42.01 | 27.90 | 8.95 × 10<sup>8</sup> |
| 1972 | 3366 | 50.56 | 1067.74 | 41.43 | 25.48 | 10.96 × 10<sup>8</sup> |
| 1976 | 3382 | 40.95 | 1238.78 | 43.73 | 28.27 | 10.68 × 10<sup>8</sup> |
| 1983 | 3378 | 51.07 | 1012.08 | 39.73 | 25.38 | 9.67 × 10<sup>8</sup> |
| 1988 | 3379 | 48.79 | 1028.70 | 40.58 | 25.29 | 7.39 × 10<sup>8</sup> |
| 1991 | 3394 | 38.17 | 1144.96 | 42.67 | 26.73 | 9.06 × 10<sup>8</sup> |
| 1999 | 33P67 | 48.28 | 1014.85 | 39.32 | 25.70 | 7.89 × 10<sup>8</sup> |
| 2007 | 33T59 | 51.82 | 1020.52 | 38.90 | 26.12 | 10.15 × 10<sup>8</sup> |
| 2011 | P1151 | 44.24 | 1113.21 | 42.03 | 25.98 | 9.41 × 10<sup>8</sup> |
| 2013 | P1197 | 42.72 | 1127.16 | 40.81 | 27.53 | 9.79 × 10<sup>8</sup> |
| 2014 | P0574 | 47.53 | 1043.57 | 39.97 | 25.98 | 9.87 × 10<sup>8</sup> |
| 2014 | P1197 | 44.75 | 1069.87 | 40.79 | 26.09 | 11.84 × 10<sup>8</sup> |
| 2015 | P1185 | 50.05 | 1066.51 | 41.02 | 25.86 | 9.90 × 10<sup>8</sup> |
| 2017 | P1366 | 39.69 | 1052.10 | 40.35 | 25.85 | 7.45 × 10<sup>8</sup> |
| 2018 | P1244 | 43.99 | 1135.09 | 42.19 | 26.73 | 8.62 × 10<sup>8</sup> |
| 2020 | P1587 | 51.07 | 1090.56 | 41.12 | 26.40 | 8.76 × 10<sup>8</sup> |
| 2020 | P1548 | 54.35 | 1008.20 | 38.86 | 25.83 | 10.03 × 10<sup>8</sup> |
| 2021 | P0953 | 41.71 | 1169.97 | 42.09 | 27.68 | 8.03 × 10<sup>8</sup> |
| 2021 | P0995 | 52.08 | 1193.15 | 43.81 | 27.13 | 8.23 × 10<sup>8</sup> |
| 2021 | P1082 | 50.31 | 1064.93 | 40.57 | 26.10 | 8.68 × 10<sup>8</sup> |
| 2021 | P1222 | 48.28 | 1054.84 | 40.63 | 25.80 | 8.73 × 10<sup>8</sup> |
| 2022 | P1413 | 47.27 | 1114.55 | 41.97 | 26.40 | 9.34 × 10<sup>8</sup> |

**Table 5.** Parameters for the Linear Regression Analysis Between Measured Stomatal Traits and Year of Hybrid Release.

|  |  |  |  |  |  |  |  |
| --- | --- | --- | --- | --- | --- | --- | --- |
|  |  | Correlation | Origin (1920 value) | Slope | R2 | p value | n |
| Total Stomatal pore area |  | -0.46 | 9.36 × 10<sup>8</sup> | -1992 | 0.21 | ≤0.05 | 27 |
| Stomatal density |  | 0.46 | 43.5 | 0.077 | 0.21 | ≤0.05 | 27 |
| Stomatal size (area) |  | -0.25 | 1057 | -0.580 | 0.06 | ns | 27 |
| Stomatal length |  | -0.18 | 39.3 | -0.010 | 0.03 | ns | 27 |
| Stomatal width |  | -0.26 | 26.7 | -0.007 | 0.06 | ns | 27 |

The table 5. presents the regression coefficients (slope), coefficient of determination (R<sup>2</sup>), and statistical significance (p-values) for the relationship between measured stomatal traits and the year of hybrid release. Positive slopes indicate an increasing trend over time, while negative slopes suggest a decline. R<sup>2</sup> values reflect the proportion of variation explained by the year of release. Significant p-values (p < 0.05) indicate a statistically meaningful relationship between the trait and hybrid release year.

**Figure 9.**
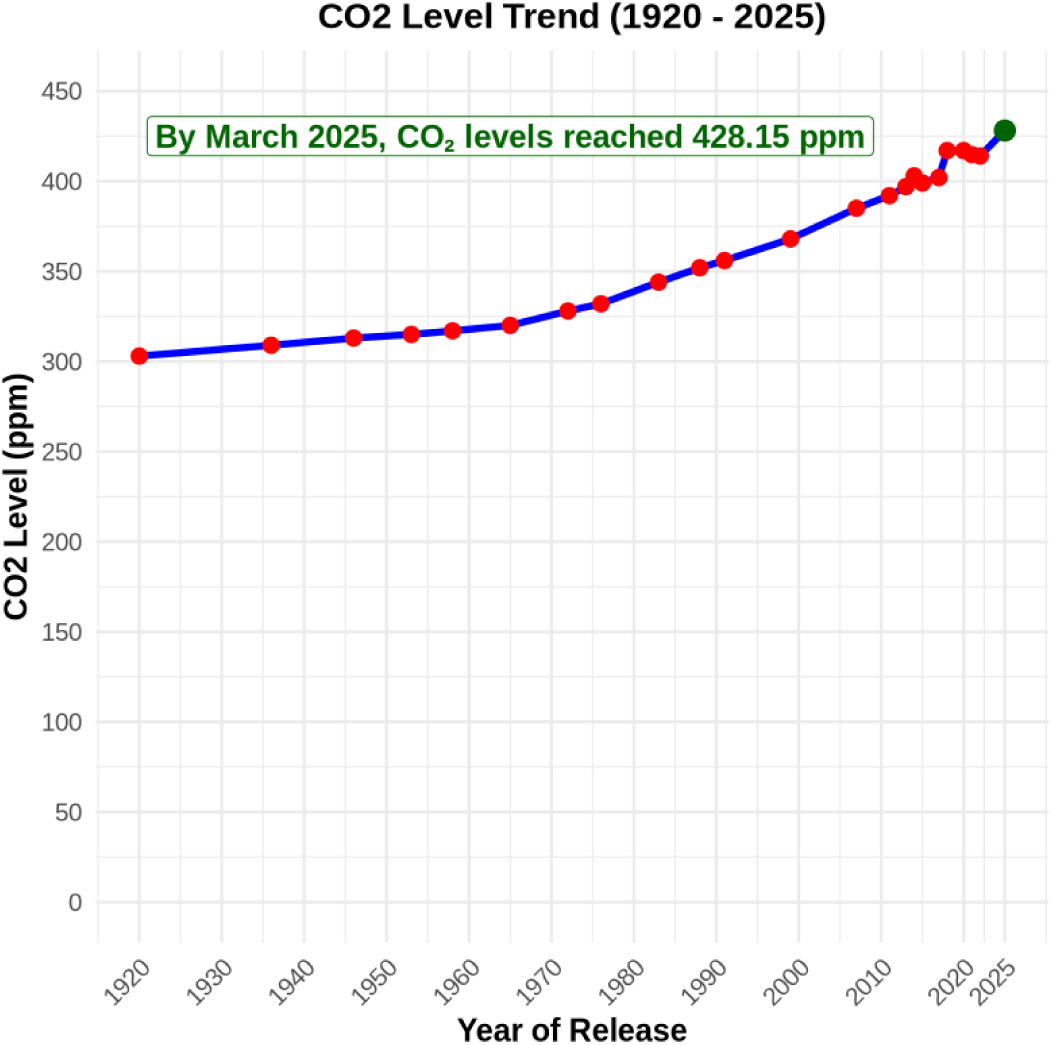
illustrates the last month of the year average carbon dioxide measured at Mauna Loa Observatory; Hawaii adapted from NOAA. The carbon dioxide data constitute the longest record of direct measurements of CO_2_ in the atmosphere on Mauna Loa. NOAA. 2025. Climate change: atmospheric carbon dioxide. Earth System Research Laboratory. Available at: www.esrl.noaa.gov. Accessed: 4/15/2025.

**Figure 10.**
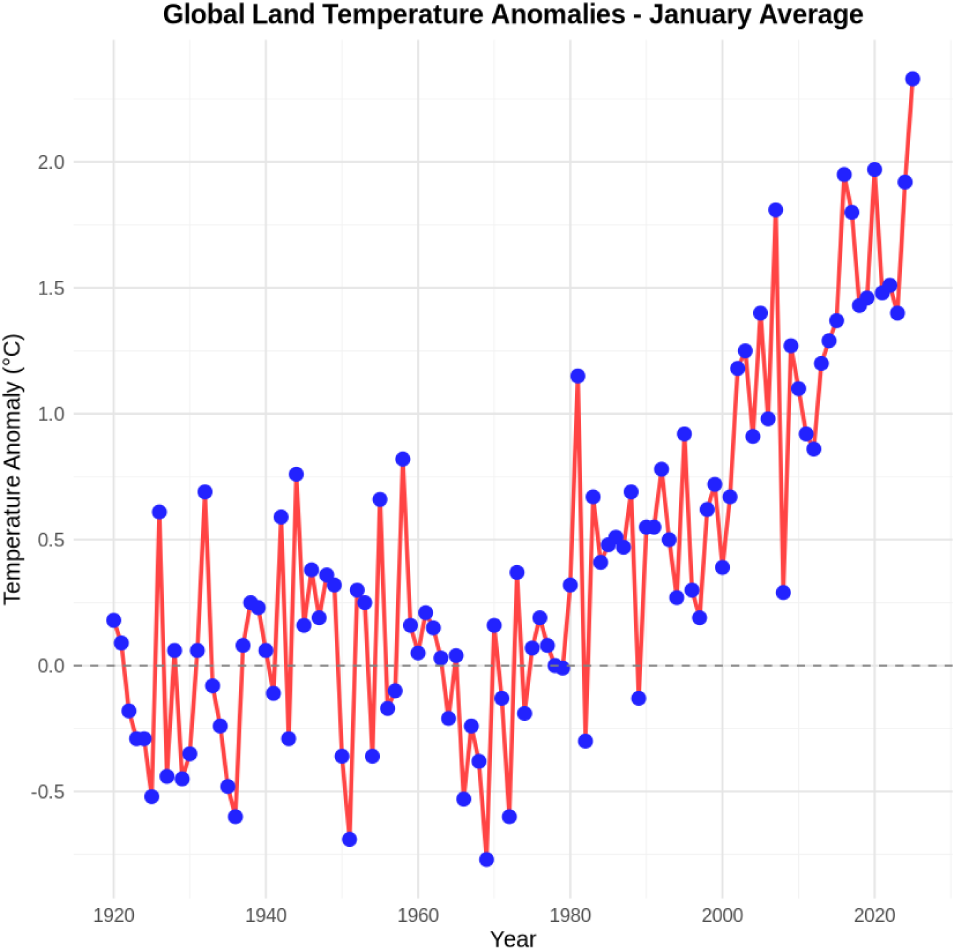
illustrates the average temperature anomaly (°C) for each January of the year, as measured by the NOAA Merged Land-Ocean Global Surface Temperature Analysis (Available at https://www.ncei.noaa.gov, accessed :4/15/2025).

**Figure 11.**
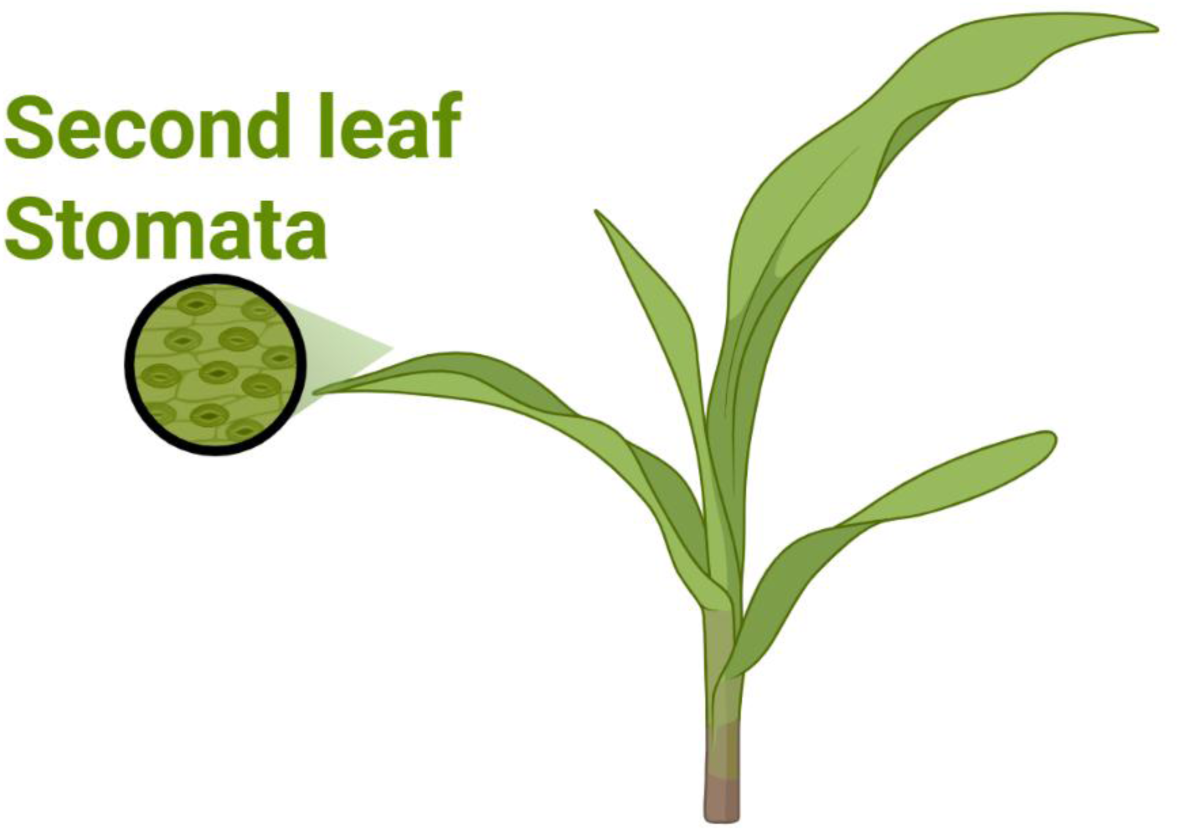
The diagram illustrates a maize plant, specifically highlighting the second leaf where stomatal traits are measured. This is the second basal leaf that emerges shortly after the first leaf during the plant’s development. An inset within the diagram shows a microscopic view of the stomata, emphasizing that measurements such as density, length, width, and area were taken from this location.

**Figure 12.**
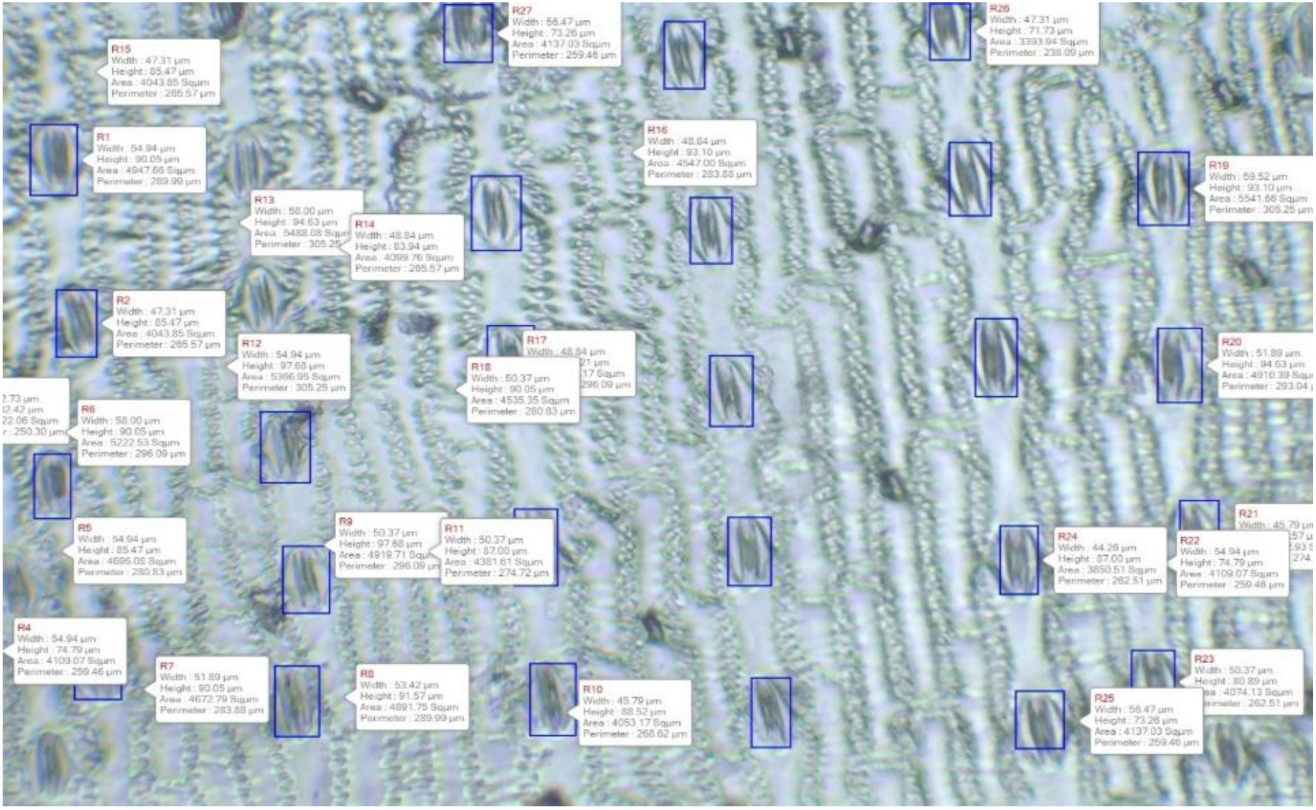
presents a microscopic view of stomatal traits, including stomatal length (μm), which measures the distance from one end of the pore to the other, and stomatal width (μm), which is the measurement perpendicular to the length (short axis). Stomatal area-size (μm^2^) is defined as the overall area of the stomatal pore. Stomatal density (stomata/mm^2^) is represented by displaying a defined surface area along with a count of the stomata within that area.

**Table 7.** Model fit (marginal and conditional R^2^) for M1 and M2, with differences (M2−M1).

| Outcome | R2 marginal M1 | R2 conditional M1 | R2 marginal M2 | R2 conditional M2 | $\Delta$ Marginal (M2-M1) | $\Delta$ Conditional (M2-M1) |
| --- | --- | --- | --- | --- | --- | --- |
| Stomatal Density | 0.080 | 0.380 | 0.080 | 0.370 | 0.000 | -0.010 |
| Stomatal Size | 0.040 | 0.670 | 0.040 | 0.670 | 0.000 | 0.000 |
| Stomatal Pore Index | 0.040 | 0.230 | 0.060 | 0.210 | 0.020 | -0.020 |
| Stomatal Length | 0.020 | 0.690 | 0.030 | 0.700 | 0.010 | 0.010 |
| Stomatal Width | 0.040 | 0.580 | 0.040 | 0.580 | 0.000 | 0.000 |
| Total Stomatal Pore Area | 0.080 | 0.360 | 0.510 | 0.570 | 0.430 | 0.210 |

**Table 8.** Computed SEM effects (unstandardized).

| Outcome | a | b | Indirect (a×b) | Direct (c') | Total Effect (c' + a×b) |
| --- | --- | --- | --- | --- | --- |
| Stomatal Density | -1.932 | -0.411 | 0.794 | 1.279 | 2.073 |
| Stomatal Size | -1.932 | -0.187 | 0.362 | -17.228 | -16.866 |
| Stomatal Pore Index | -1.932 | -475.643 | 919.068 | 709.880 | 1628.948 |
| Total Stomatal Pore Area | -1.932 | 39.8M | -76.9M | 14.7M | -62.2M |

*Total stomatal pore area, SD×SS scaled to mm<sup>2</sup>.*

**Table 5.**
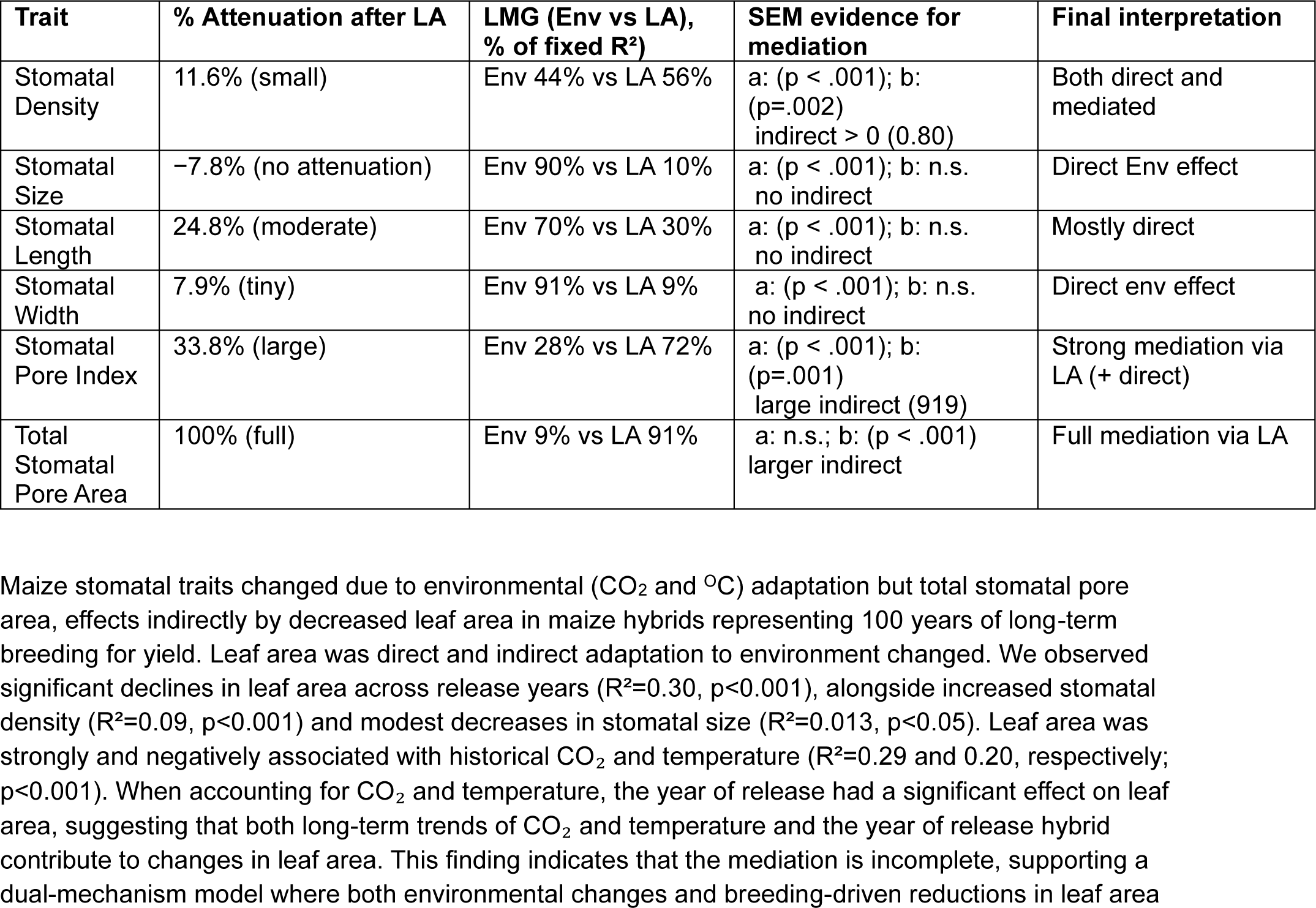
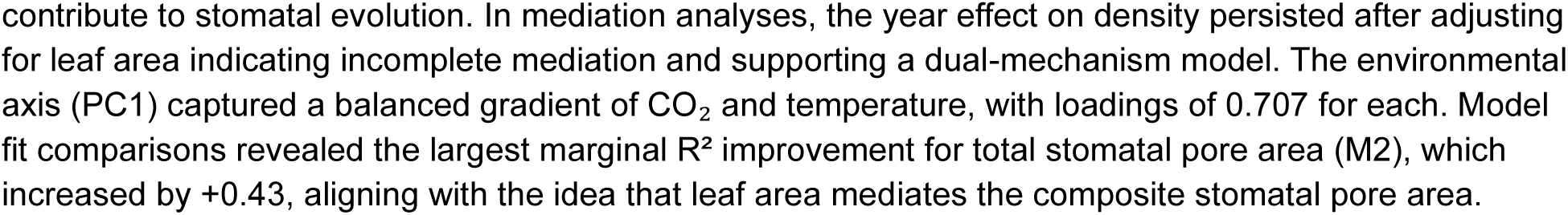
Attenuation of zEnv, LMG fixed-effects contributions, and mediation verdicts.

**Figure 13.**
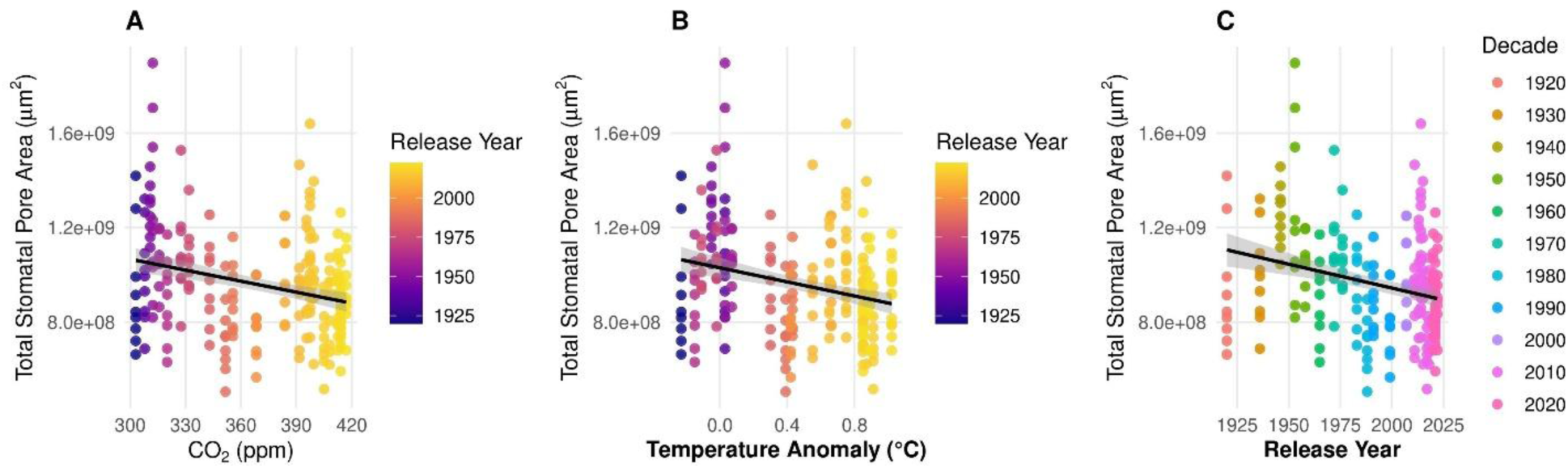
Total stomatal pore area versus (A) atmospheric CO_2_ (ppm), (B) mean temperature anomaly (°C), and (C) release year. Points are replicate observations colored by release year (A–B) or decade (C). Lines are linear regressions with 95% confidence bands.

**Figure 14.**
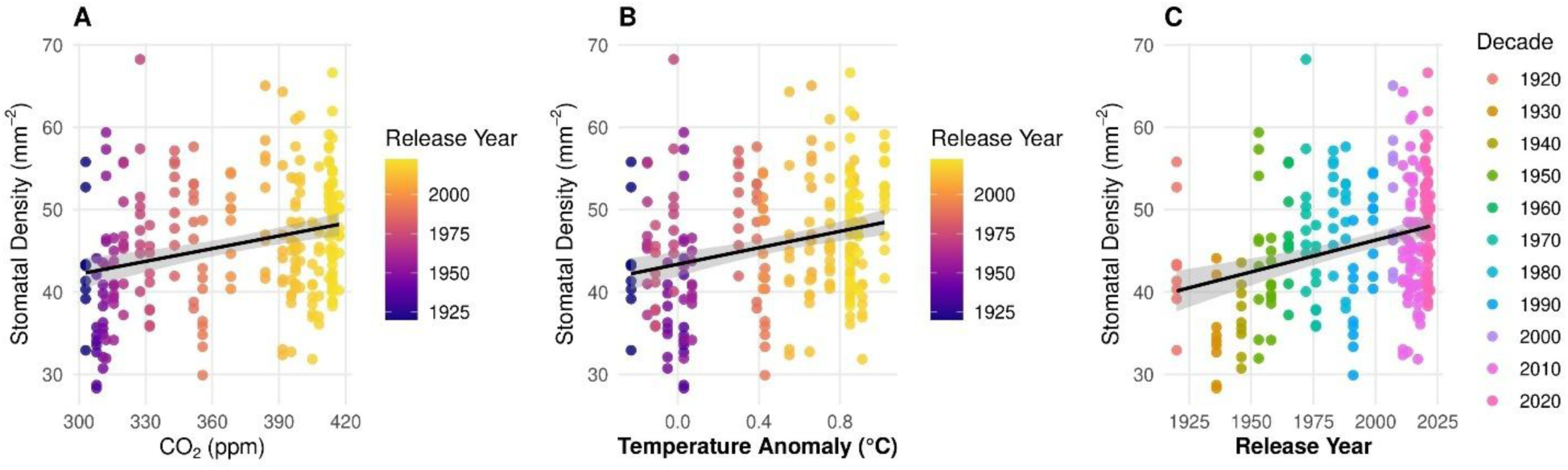
Stomatal density versus (A) atmospheric CO_2_ (ppm), (B) mean temperature anomaly (°C), and (C) release year. Points are replicate observations colored by release year (A–B) or decade (C). Lines are linear regressions with 95% confidence bands.

**Figure 15.**
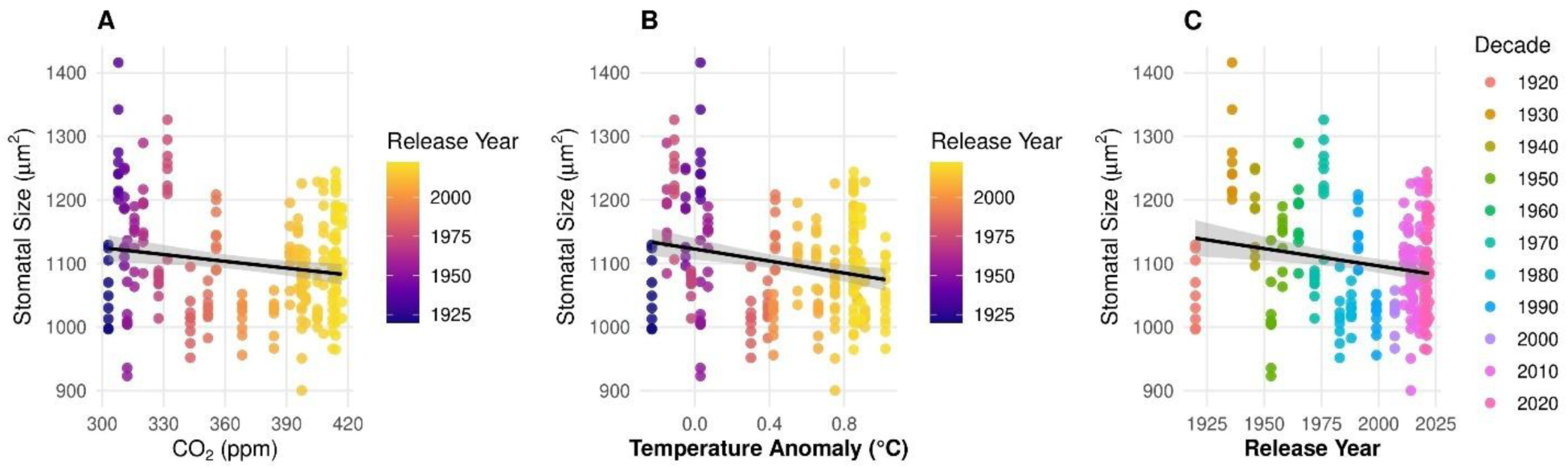
Stomatal size versus (A) atmospheric CO_2_ (ppm), (B) mean temperature anomaly (°C), and (C) release year. Points are replicate observations colored by release year (A–B) or decade (C). Lines are linear regressions with 95% confidence bands.

**Figure 16.**
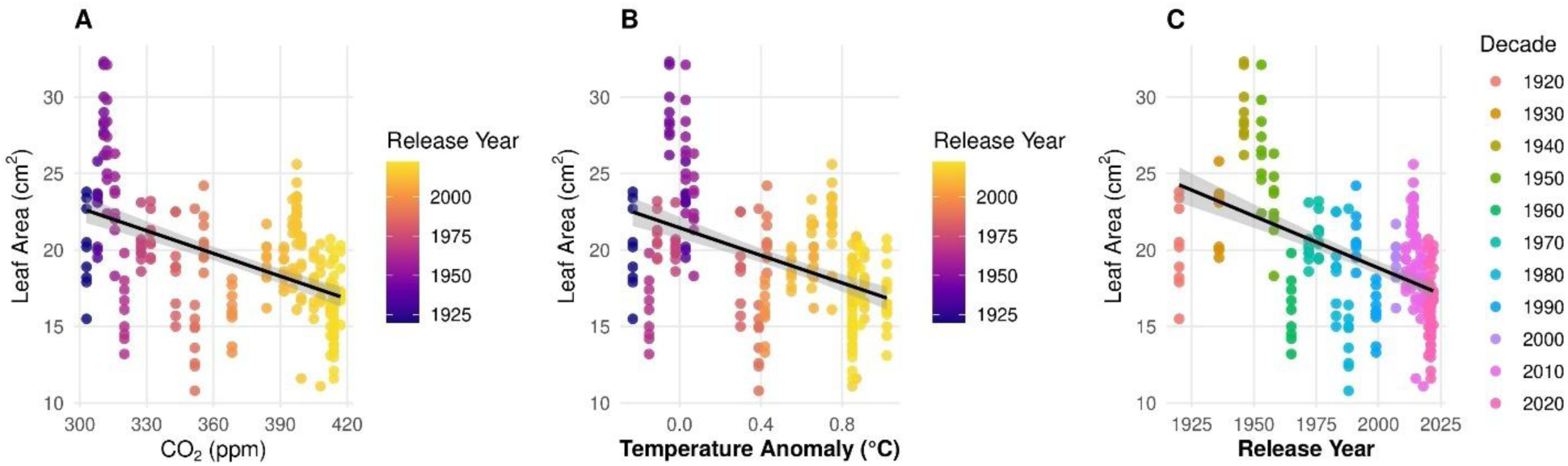
Leaf area versus (A) atmospheric CO_2_ (ppm), (B) mean temperature anomaly (°C), and (C) release year. Points are replicate observations colored by release year (A–B) or decade (C). Lines are linear regressions with 95% confidence bands.

**Figure 17.**
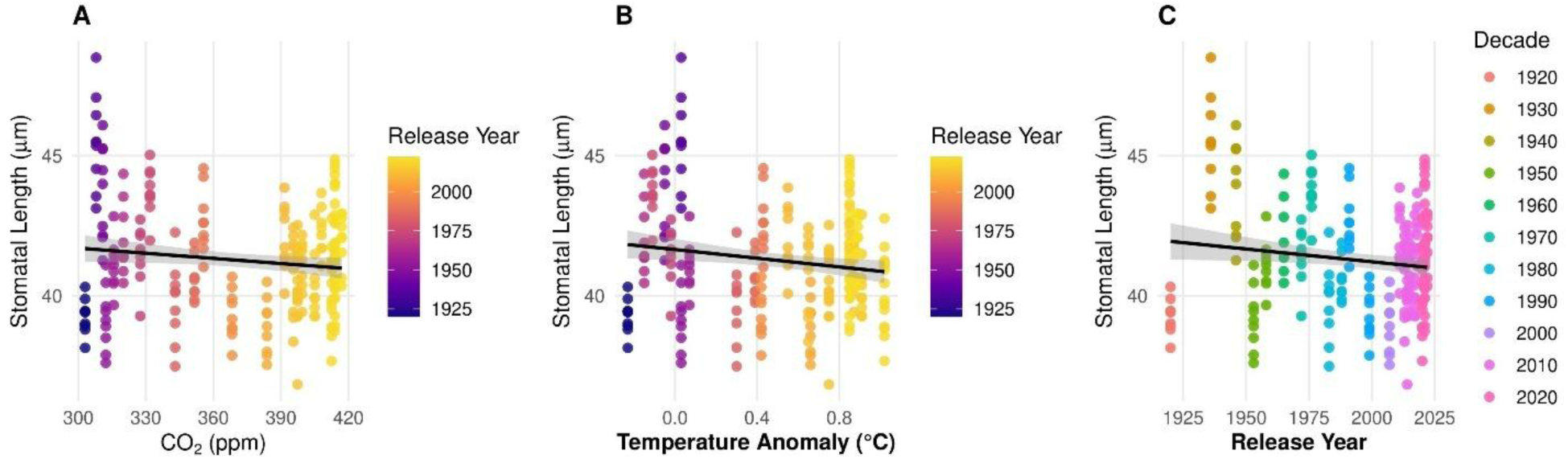
Stomatal length versus (A) atmospheric CO_2_ (ppm), (B) mean temperature anomaly (°C), and (C) release year. Points are replicate observations colored by release year (A–B) or decade (C). Lines are linear regressions with 95% confidence bands.

**Figure 18.**
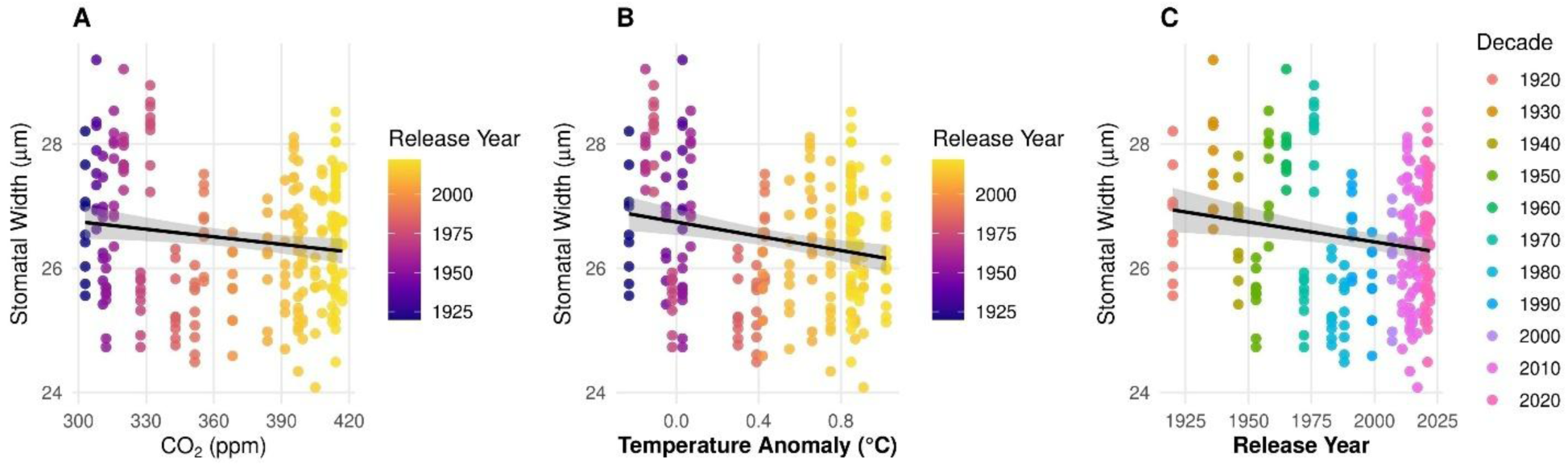
Stomatal width versus (A) atmospheric CO_2_ (ppm), (B) mean temperature anomaly (°C), and (C) release year. Points are replicate observations colored by release year (A–B) or decade (C). Lines are linear regressions with 95% confidence bands.

**Figure 19.**
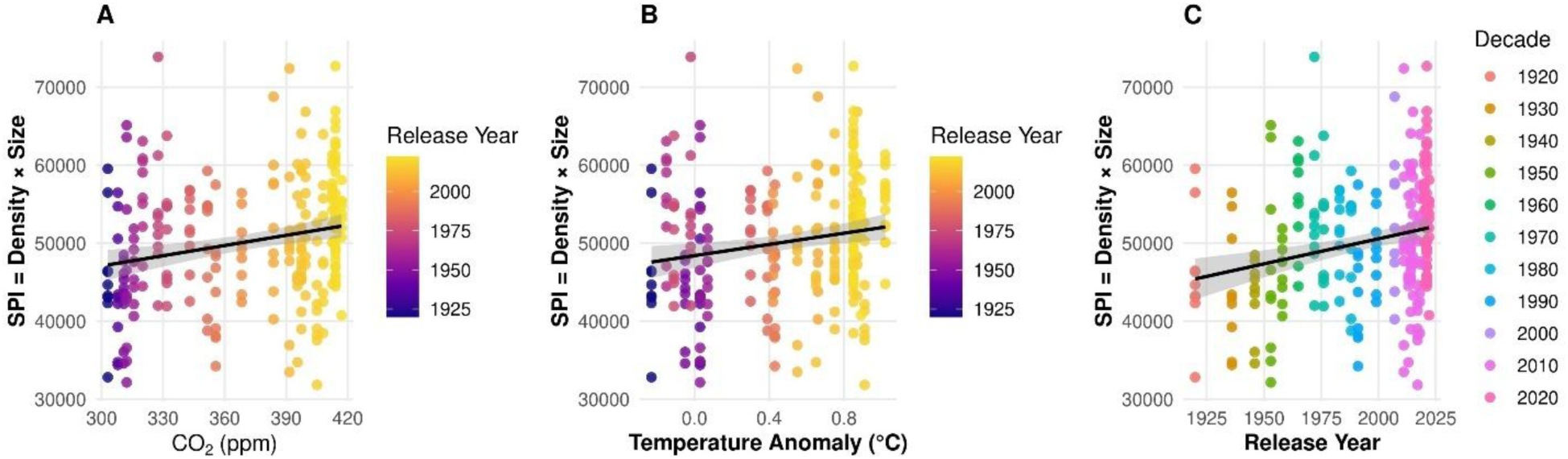
Stomatal density x size versus (A) atmospheric CO_2_ (ppm), (B) mean temperature anomaly (°C), and (C) release year. Points are replicate observations colored by release year (A–B) or decade (C). Lines are linear regressions with 95% confidence bands.

